# Dendritic turnover zones support switching between rules without expanding the cortical engram

**DOI:** 10.64898/2026.08.26.747194

**Authors:** Ioanna Pandi, Spyridon Chavlis, Hatem Oraby, Mostafa A. Nashaat, Matthew E. Larkum, Athanasia Papoutsi, Panayiota Poirazi

## Abstract

Flexible behavior depends on frontal circuits that use experience to adjust subsequent decisions. Yet how synaptic connections in these circuits are structurally reorganized when new associations compete with established ones remains unclear. Here, we used a novel rule-switching behavioral paradigm, longitudinal spine imaging, and circuit modeling to investigate how dendrites accommodate changing associations. Inactivation of secondary motor cortex (M2) impaired rule switching but not learning or maintenance in control conditions. Switching increased spine turnover on apical tuft dendrites in M2, concentrating spine formation, elimination, and new-spine clustering within persistent dendritic turnover zones, with pre-existing spines lost early during adaptation. The model reproduced these dynamics and predicted preferential elimination of synapses encoding the prior rule. Preventing dendritic compartment reuse increased the neuronal population representing the rules. These findings suggest that persistent turnover zones support synaptic reorganization during behavioral updating, with reuse offering a potential economy in neuronal recruitment.

## Introduction

Learning from experience allows us to adapt our behavior as the world changes. The brain must build on existing knowledge but also revise associations that no longer guide appropriate behavior, a central challenge for artificial systems^1^. Dendritic spines are the principal structural units across which adaptive learning is implemented and are therefore central to understanding behavioral adaptation^2,3^. Longitudinal two-photon imaging has documented their formation, stabilization, and elimination during learning across brain areas^4–9^. A consistent organizing principle has emerged from these studies: learning-related spines are not distributed uniformly along the dendrite but cluster within small neighborhoods, where they integrate with previously potentiated synapses and are recruited in a branch-specific manner^4,5,10–13^. Converging evidence identifies short dendritic segments, often on the order of 10 µm, as sites of coordinated synaptic activity and local interactions between plasticity mechanisms^14–18^. This suggests that neighboring synapses can form local functional clusters whose properties depend on their organization as well as their individual strengths. Such clustering amplifies the impact of correlated inputs on dendritic spiking and somatic output^19–21^, is engram-specific in motor circuits^22,23^, and is increasingly viewed as the structural basis of memory storage^24–27^. Critically, optogenetic shrinkage of a learning-related cluster disrupts the corresponding memory, establishing these domains as causal substrates of a specific memory^28^.

Despite the abundance of evidence that the brain is constantly reconfiguring synaptic inputs in the structural sense, it remains unclear how the formation and elimination of synapses is coordinated within dendrites when new learning competes with an established association. At the level of neuronal ensembles, related experiences are thought to be allocated to overlapping populations whereas unrelated ones are kept apart^29–31^. Does the same principle govern where new synapses form within the dendritic tree? According to Cichon and Gan^10^, two independent motor skills recruit coordinated spine plasticity on largely non-overlapping branches in the motor cortex, supporting the view that distinct episodes can be physically segregated to minimize interference. Conversely, Sehgal and colleagues demonstrated that the same dendritic domains are co-allocated to spine clusters from two contextual memories formed close in time, establishing that domain sharing can link memories across time^13^, a possibility also anticipated by computational work^27^. Cortical pyramidal neurons can therefore either segregate or reuse their dendritic branches. However, what determines whether successive learning episodes recruit shared or separate dendritic domains remains unclear. One possibility is that this organization depends on the relationship between the associations being learned. We therefore ask how synapses are reorganized when a new association competes with an established one.

When a previously rewarded contingency must be replaced by a new one, the circuit must simultaneously acquire a novel association and disengage from the prior strategy. What happens to the synaptic configuration encoding the prior contingency remains uncharacterized. Prior work has shown that the spines formed during fear conditioning are preferentially eliminated during subsequent extinction^6^, suggesting that contingency-specific synapse elimination is itself a structurally regulated process. Whether comparable dynamics target the dendritic compartments engaged by an old rule when a new one becomes relevant, and whether such elimination is spatially coupled to the formation of new clustered spines, has not been tested. The nature of the reorganization is equally unsettled. Functional imaging of apical tuft dendrites in anterolateral motor cortex has shown that tuft activity is selectively reorganized during rule-switching, with active synapses changing their spatial configuration^32^. Synaptic activity was more spatially clustered during performance of the more difficult rule. However, such effects could arise from the different activity patterns of presynaptic neurons, presynaptic axonal rewiring, or from changes in synaptic weight and inhibitory gating that leave postsynaptic connectivity intact. The longitudinal structural imaging needed to distinguish these possibilities across a rule transition has not been performed.

How are synaptic connections structurally reorganized during learning? And, what computational advantage might this organization provide? The answer to the first question requires experimental data from longitudinal spine imaging. However, the second requires a theoretical model in order to test the contribution of reorganization at the network level, using different regimes of dendritic reuse and segregation. The model also allows for the quantification of representational efficiency (engram economy) and coding capacity of alternative regimes^33^, connecting cellular and biophysical mechanisms to systems and cognitive functions. Conversely, abstract reinforcement-learning and rate-based models that capture flexible behavior at the population level^34–37^, operate above the level at which structural plasticity acts and cannot represent the physical clustering of synapses.

We address these two fundamental questions in the rodent secondary motor cortex (M2). M2 integrates sensory information with reward outcomes^38–40^, exhibits choice-predictive activity that precedes movement^41^, and is causally required for updating action plans under conflicting cues^39,42,43^. The apical tuft dendrites of layer 5 pyramidal neurons, the major output of M2, receive long-range inputs from orbitofrontal cortex, sensory areas, and higher-order thalamus^44,45^, i.e. the contextual signals that adaptive behavior requires^46,47^. We used longitudinal two-photon imaging to detect where spines appear and disappear and used this data in a biologically grounded spiking network model in which multicompartmental pyramidal neurons undergo calcium-dependent plasticity and structural turnover to reveal the rule identity of an eliminated synapse, and determine whether the spatial structure of plasticity causally constrains circuit-level adaptation.

We found that M2 is selectively required for resolving competition between an established and an emerging rule, but not for rule learning or maintenance. This adaptation is accompanied by spatially confined dendritic “turnover zones” that host both clustered formation of new spines and elimination of new and pre-existing spines. Within these dendritic domains, pre-existing spines are eliminated preferentially early after the switch, whereas newly formed spines continue to undergo turnover throughout adaptation. Our network model reproduces key features of these structural dynamics and predicts that the early-eliminated spines preferentially encode the prior rule. Preventing dendritic reuse in the model increases the number of neurons recruited to represent the rules, suggesting a computational advantage of having shared dendritic domains. Together, these findings, support a view of local dendritic domains as sites where coordinated synapse formation and elimination reorganize existing connections during behavioral updating.

## Results

### Behavioral adaptation under competitive demand requires M2

To investigate the structural plasticity that supports flexible behavior under competitive demand, we first developed a novel cross-modal rule-switch task for head-fixed mice (**Fig. 1a, b**, **Extended Data Fig. S1**). Mice were trained to respond to auditory or visual cues by rotating an air-floating platform toward one of two choice ports, a configuration that allows active behavioral choice under head fixation^42,48^. Mice were first trained on a sound discrimination rule (SR): an up-sweep tone (from 11 to 15 kHz) indicated reward at the right port, a down-sweep tone (from 16 to 12 kHz) at the left (**Fig. 1a**). Trials were self-initiated by entering the central port; correct choices delivered a 5.5 µL water reward, while incorrect choices triggered a punishment tone and time-out. Mice were considered experts when performance exceeded 75% correct responses for two consecutive sessions.

**Fig. 1.**
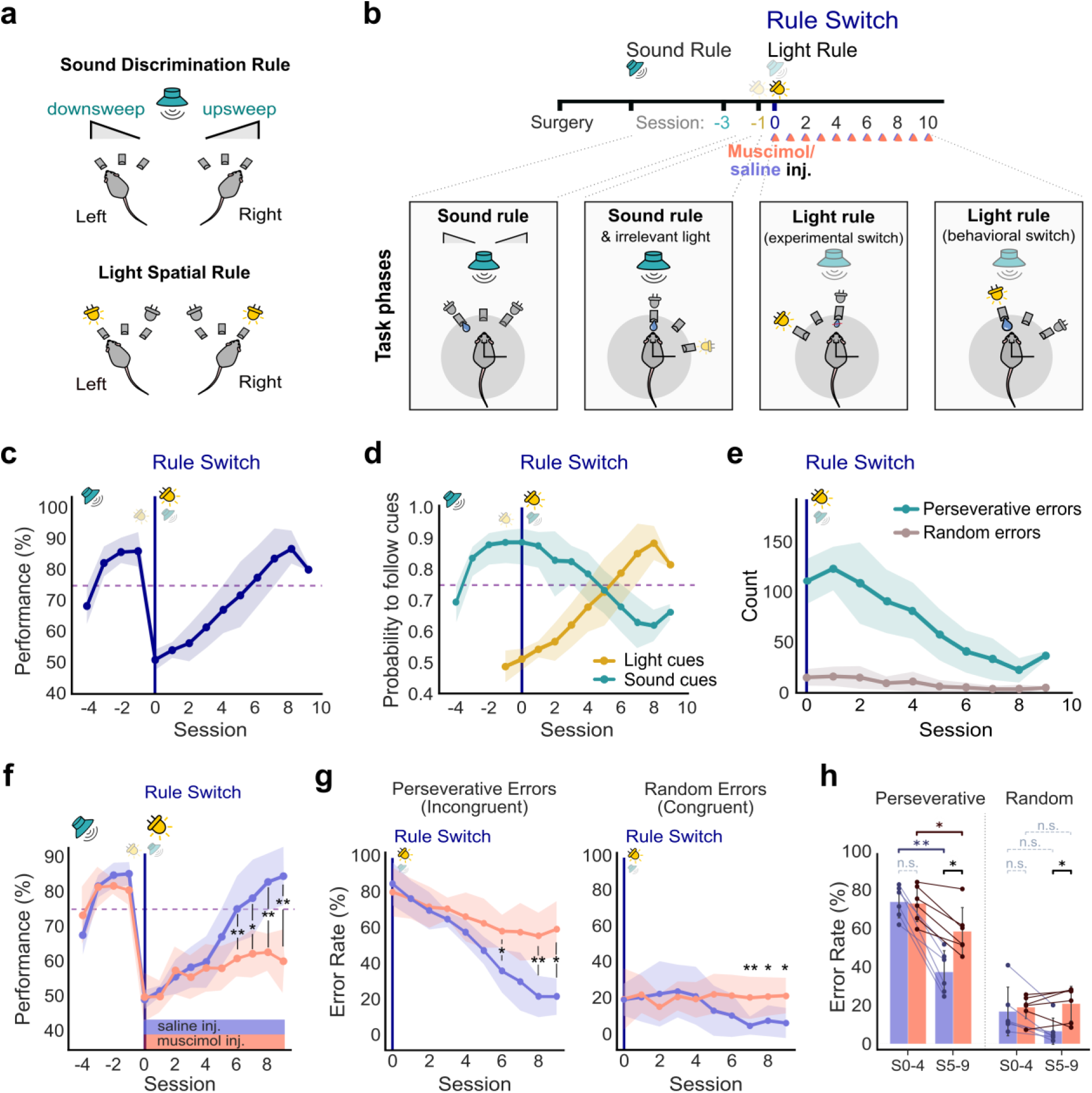
M2 is selectively required for cross-modal rule-switching, but not for rule maintenance, in a novel head-fixed task. **a**, Task design. In the sound rule (SR), an up-sweep tone (from 11 to 15 kHz) indicated reward at the right port and a down-sweep tone (from 16 to 12 kHz) at the left. In the light rule (LR), the illuminated choice port indicated reward and sound cues were no longer rewarded. **b**, Training paradigm. Mice were first trained on the SR (criterion: >75% correct over two consecutive sessions). A non-rewarded distractor light cue was introduced for one session (Session -1), followed by the rule switch (Session 0) and subsequent LR acquisition. For the results shown in panels **f, g**, muscimol or saline injections (triangles) were administered bilaterally into M2 prior to each session from Session 0 onward in the inactivation cohort. **c**, Performance across training sessions (% correct trials) for the imaging cohort (n=11 mice). Performance drops to chance at Session 0 and recovers above the 75% criterion (dashed line) within ∼6 sessions. **d**, Probability of choices consistent with the sound cue (turquoise) or the light cue (yellow) across sessions. Following the switch, mice initially continue to follow sound cues for several sessions (n=11 mice). **e**, Error classification. Perseverative errors (choices consistent with the previously rewarded SR; teal) dominated the early post-switch period and declined with learning. Random errors (choices inconsistent with both rules; gray) remained low throughout the learning sessions (n=11 mice). **f,** M2 inactivation impairs rule-switching. Performance for saline- (blue, n=6) and muscimol-treated (orange, n=6) mice; muscimol-treated mice failed to reach criterion across sessions 0-10 (mixed ANOVA, F=3.78, p=0.0006; per-session Mann–Whitney U test, sessions 6-9, *p<0.05, **p<0.01). **g**, Right: Normalized error rates for perseverative (% of incongruent trials) errors. M2 inactivation caused an increase in perseverative errors (Saline group: 6 mice, Muscimol group: 6 mice, 117 total sessions, Mann-Whitney U test, sessions 6 and 9: p<0.05, session 8: p<0.01). Left: Normalized error rates for random (% of congruent trials) errors. M2 inactivation caused an increase in random errors (Saline group: 6 mice, Muscimol group: 6 mice, Random errors: session 7, p<0.01 and sessions 8 and 9: p<0.5, Mann-Whitney U test). **h**, Normalized error rates for perseverative and random errors across early (Sessions 0–4) and late (Sessions 5–9) phases for Saline (blue) and Muscimol (orange) cohorts. Shaded bars and whiskers represent cohort mean ± SD, overlaid with connected lines tracing individual mouse trajectories across phases. Between-cohort differences and longitudinal within-cohort extinction were assessed using Welch’s two-sample t-test and two-tailed paired t-tests, respectively, with Bonferroni correction for multiple comparisons. Significance is denoted as *p < 0.05 and n.s. (p > 0.05), with the vertical dashed line separating error types. Shaded areas indicate ±1 standard deviation (SD). Full statistics and sample sizes for all comparisons are provided in **Supplementary Table 1**.

To test whether mice could update their strategy against a competing cue, we introduced a rule switch (**Fig. 1b**). A non-rewarded light cue that was co-presented with the auditory stimulus for a single session (**Fig. 1b** top, Session -1) left performance unchanged, consistent with continued use of the auditory rule (**Fig. 1c**). At Session 0 the rewarded modality changed: the illuminated port now signaled reward (light rule, LR), while the still-present sound cues became irrelevant. Mice thus had to suppress the previously rewarded auditory strategy and redirect their attention to the visual modality, which is the defining feature of a competitive switch. Performance fell to chance and recovered to expert levels over 6-7 sessions (**Fig. 1c**; n=11 mice). Early adaptation was dominated by perseverative errors (errors on incongruent trials, where choices followed the former SR) which declined as learning progressed. Random errors (errors on congruent trials, consistent with neither rule) remained low throughout the adaptation period (**Fig. 1d, e**). The predominance of perseverative errors indicates that the delay in LR acquisition reflected persistence of the prior strategy rather than a failure to detect the new cues; choices showed no side bias (**Extended Data Fig. S1**).

We next asked whether M2 is causally required for this adaptive behavior. Bilateral infusion of muscimol, a GABA_A_ receptor agonist, before each post-switch session (**Fig. 1b**) severely impaired rule-switching, as the inactivated mice failed to reach the expert performance criterion within the imaging window (**Fig. 1f**; saline n=6, muscimol n=6; mixed ANOVA, F=3.78, p=0.0006), consistent with prior reports^39,43,49^. Muscimol inactivation elevated perseverative errors (calculated as the error rate on incongruent trials; **Fig. 1g**, **right**; sessions 6, 8, and 9, Mann-Whitney U test, p < 0.05), indicating a prolonged commitment to the prior strategy. Normalized random errors (calculated on congruent trials) were also significantly elevated relative to saline controls in later sessions (**Fig. 1g**, **left**; sessions 7-9, Mann-Whitney U test, p < 0.05), indicating that mice with M2 inactivation additionally failed to learn the new contingency. In the late phases of learning (sessions 5-9), the saline group resolved both error types to successfully switch to the new rule, whereas the muscimol group remained impaired on both perseverative (58.2% vs. 37.2%, p < 0.05) and random errors (20.8% vs. 6.5%, p < 0.05; Fig. 1h). These results indicate persistent use of the prior strategy together with impaired acquisition of the new contingency when M2 is inhibited. Movement and sampling times were unaffected (**Extended Data Fig. S2d**), arguing against a gross impairment of movement execution^50^.

Two control experiments characterized the limits of this requirement. When the rule changed mid-session (Session 0, at trial ∼60), inactivated and control mice showed no significant difference in SR accuracy before the switch (**Extended Data Fig. S2b**; Mann–Whitney U test, U=17.0, p=0.93); the absence of a significant difference is consistent with M2 not being necessary for SR execution. When mice learned the LR without competing sound cues, we detected no significant effect of inactivation on acquisition (**Extended Data Fig. S2e**; mixed ANOVA, p=0.46; n=3 per group). Taken together, the inactivation data indicate that M2 is preferentially required for resolving the competition between an established and an emerging rule, with limited contributions under conditions in which conflict is absent. This preferential causal role identifies M2 as the substrate in which to ask how structural plasticity supports behavioral adaptation under competitive environmental demands.

### Rule-switching elevates spine turnover dynamics on M2 apical tuft dendrites

Having established M2 as a key region for adaptation under competitive demand, we asked whether the rule switch is accompanied by structural reorganization of its synaptic inputs. We performed longitudinal *in vivo* two-photon imaging of apical tuft dendrites of M2 layer 5 pyramidal neurons in Thy1-GFP-M mice, in which a sparse subset of these neurons expresses GFP at densities suitable for resolving individual spines (**Fig. 2a**;^51^. The same dendritic segments (∼50 µm) were tracked at ∼2-day intervals across the rule-switch period, and spine gain and loss were quantified from co-registered z-stacks of each segment (**Fig. 2c, d**). As imaged dendritic segments were not explicitly traced to their respective neuronal bodies, the analyzed data represent longitudinally tracked samples of the apical tuft arbor rather than reconstructions of complete tufts.

**Fig. 2.**
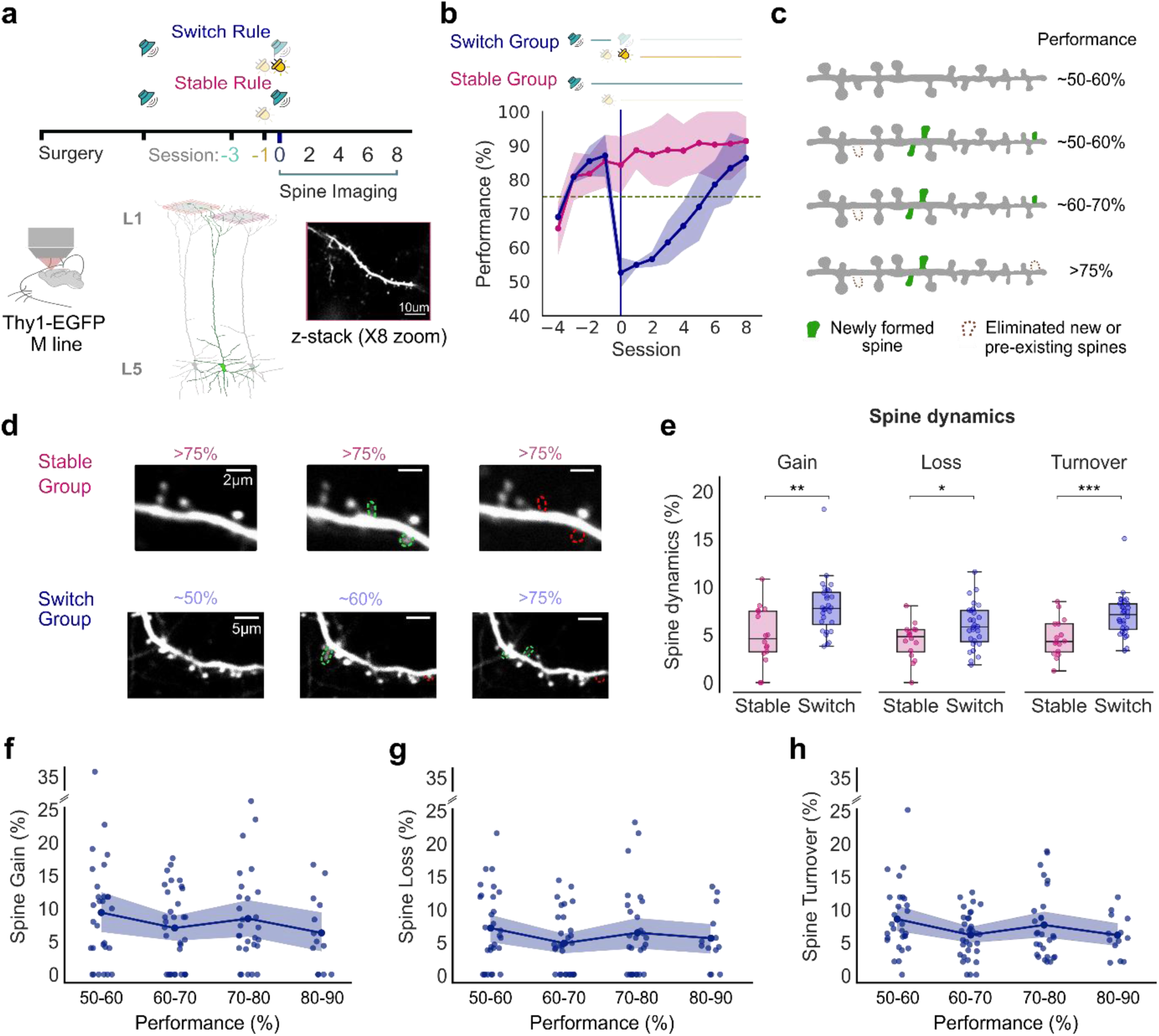
Rule-switching elevates spine turnover. **a**, Experimental design. Apical tuft dendrites of M2 layer 5 pyramidal neurons were imaged longitudinally in Thy1-GFP-M mice using two-photon microscopy. Same-segment imaging was performed every ∼2 days from Session 0 (Rule Switch) onwards, in both Switch and Stable Rule groups. Cranial windows were placed over the M2 (AP +1.8 mm, ML +0.75 mm) and z-stacks of identified dendritic segments (∼50 µm) were acquired at 8× zoom. **b**, Behavioral performance across imaging sessions for Switch (dark purple, n=4 mice) and Stable (magenta, n=4 mice) Rule groups. Threshold performance: 75% correct (dashed line). Shaded areas indicate ±1 SD. **c**, Schematic of spine classification. Spines present in one imaging session and absent in the previous were classified as formation events (left); those present in one session and absent in the subsequent were classified as elimination events (right). Classification was performed on co-registered z-stacks of the same segment. **d**, Representative two-photon images of dendritic segments from Stable (top) and Switch (bottom) groups across three sessions. Green arrowheads: newly formed spines. Red arrowheads: eliminated spines. **e,** Spine gain, loss and turnover expressed as a percentage of total spines per dendrite, averaged across the imaging window. The Switch group exhibited significantly elevated dynamics across all three metrics (Switch: n=4 mice, 28 dendrites, 645 spines; Stable: n=4 mice, 15 dendrites, 358 spines; gain: p=0.0022, Mann–Whitney U test; loss: p=0.028, t-test; turnover: p=0.0007, Mann–Whitney U test; **p<0.01, ***p<0.001). **f, g, h** Spine gain, loss and turnover, respectively, as a function of performance across the adaptation period (Switch: n=4 mice, 28 dendrites, 103 data points in each condition: spine gain, spine loss and spine turnover. Detailed statistics, p values, and sample sizes for all comparisons are provided in **Supplementary Table 1**.

To dissociate the structural correlates of flexible learning from those of platform navigation or initial rule acquisition, we also imaged a within-task control group. Stable Rule mice were trained on the SR to the same expert criterion as the Switch group and then continued to perform the SR for an equivalent number of sessions, with the light cue present but unrewarded (**Fig. 2b**). This design matches sensory exposure, motor demands, and imaging schedule across groups, isolating the rule-switch demand as the only distinguishing variable; imaging began on the first day of the SR in both groups (session 0). Baseline dendritic architecture was equivalent, because the spine density and imaged dendritic length did not differ at Session 0 (**Extended Data Fig. S3a, b**; length, Mann–Whitney U test, p=0.60; density, t-test, p=0.84).

Rule-switching markedly elevated structural dynamics. Spine gain, loss, and overall turnover rates were all higher in the Switch than in the Stable group across the imaging window (**Fig. 2e**; Switch, n=4 mice, 28 dendrites, 645 spines; Stable, n=4 mice, 15 dendrites, 358 spines; gain, p=0.0022; turnover, p=0.0007, Mann–Whitney U test; loss, p=0.028, t-test) and remained elevated throughout the adaptation period (**Fig. 2f-h**). Because the elevation reflected both the formation of new spines and the elimination of pre-existing ones, the switch drives bidirectional reorganization rather than a unidirectional gain or loss of inputs. Conversely, the Stable group showed lower dynamics despite identical sensory cues and motor demands, indicating that the elevated remodeling tracks the cognitive demand of updating an established strategy rather than ongoing perceptual or motor engagement. Spine density in the Switch group nonetheless increased modestly relative to baseline (**Extended Data Fig. S3c**; Pearson r=0.49, p=0.04), indicating that gain slightly exceeded loss, a finding consistent with the transient net expansion reported during the early active phase of learning in motor and sensory cortices^8,9,52^.

### Spine remodeling is spatially concentrated within stable dendritic turnover zones

Given the marked remodeling that accompanied flexible learning, we next asked whether spine turnover is distributed uniformly along the dendrite or concentrated at particular sites. We mapped the precise location of each spine event (gain or loss) along each dendritic segment throughout the switch-period and computed the local event density using sliding 5-µm windows (see Methods). This allowed us to identify locations where the number of turnover events peaked. Turnover zones were defined as dendritic regions of high turnover, in which density peaks exceeded the 90th percentile of local-density threshold, and verified with Monte Carlo permutation analysis (see Methods, **Fig. 3a**). Turnover zones were highly prevalent in our imaging dataset: 21 of the 28 dendritic segments in the Switch group contained at least one turnover zone, while 6 dendrites contained two (**Fig. 3b**). Compared to the null model in which event positions were shuffled across biologically valid spine locations on the same segment, both the number of turnover zones per segment and their peak amplitudes were significantly elevated (**Fig. 3c, d**; p<0.0005, Mann–Whitney U test), and in segments with multiple zones adjacent peaks lay on average 24 µm apart (range: 18-30 µm, **Fig. 3e**). An equivalent, above-chance concentration of turnover events was also present in the Stable group (**Extended Data Fig. S4**), indicating that these zones are a general organizational feature of M2 L5 tuft dendrites.

**Fig. 3.**
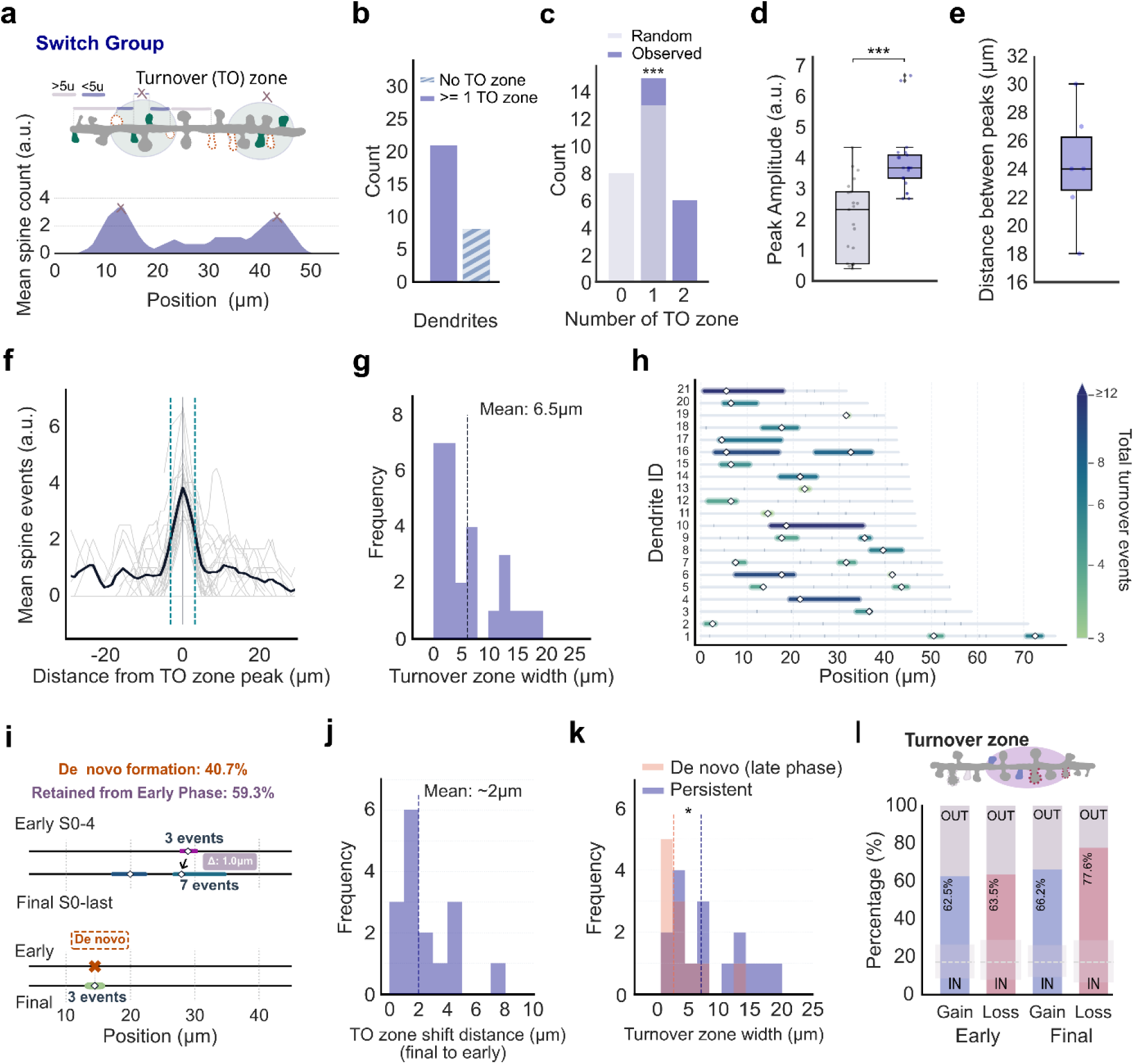
Spine remodeling is non-uniform and concentrates within spatially stable dendritic turnover zones. **a**, Example Switch-group segment showing the spatial distribution of spine turnover events. Mean spine-event count was computed in sliding 5-µm bins along the segment; peaks exceeding the 90th percentile local-density threshold (2.7 events for the Switch group) and separated by at least 10 µm define turnover zones. Two zones are identified in this example. **b**, Distribution of turnover zones per dendritic segment in the Switch group; 21 of 28 segments contained at least one turnover zone (4 mice, 28 dendrites). **c**, Turnover zone count per segment in observed data versus a shuffled null model in which event positions were redistributed across the experimentally-identified spine locations on the same segment (observed: 4 mice, 21 dendrites, 244 events; null: 1,000 shuffles per segment; ***p<0.0005, Mann–Whitney U test). **d**, Turnover zone peak amplitude (mean spine-event count at peak) in observed versus null data (***p<0.0005, Mann–Whitney U test). **e**, Distance between adjacent zone peaks in segments containing two or more turnover zones (mean=23.8µm). Box plots show median, interquartile range and minimum/maximum. **f**, Method for estimating zone width. Thick black line: mean spine-event-count curve averaged across all zone-containing segments; grey lines: individual segment curves; vertical cyan dashed lines mark the turnover zone boundaries, defined by the physical span of contiguous clustered events (inter-event distance <5μm. **g**, Distribution of turnover zone widths, mean = 6.5 µm. **h**, Positioning of turnover zones for the 21 dendritic segments containing them. The position is relative to the start and end of the imaged segment, indicated by the horizontal gray lines; color encodes the total number of turnover events (spine gain or loss) that took place within the turnover zone, measured over the entire adaptation period**. i**, Turnover zone persistence. Zones were detected from events in the early phase (sessions 0–4) and from all events across the adaptation period (sessions 0–last). Most zones present at the end of learning (59.3%) were already detectable in the early sessions (0-4) following the switch; the remainder (40.7%) emerged only once later events were included. **j**, Turnover zone stability: distribution of the shift in zone-center position between the early and final phases, ranging from 0 to 8 µm (mean ≈ 2 µm), indicating that persistent zones are spatially stable across adaptation. **k**, Distribution of widths of zones emerging after the early phase (median = 2.4 µm) and of zones present in both analyses (median = 6.8 μm). Later-emerging zones exhibit significantly narrower spatial domains than persistent zones (p = 0.0146, Mann-Whitney U test). **l**, Localization of formation and elimination events relative to zones (IN: inside vs. OUT: outside) across the two phases of the adaptation period. In all groups, the fraction of events within turnover zones exceeded chance levels, and within-zone elimination events were higher in the final than in the early phase (77.6% vs. 63.5%, respectively). Dashed lines indicate chance levels (Monte Carlo permutations) for all groups: mean values are ∼17% for all groups with range: [8-26%] for early phase formation events, [8-29%] for early phase elimination events, [9-26%] for the final phase formation events, and [6-29%] for the final phase elimination events (one-sided Wilcoxon signed-rank test, all groups: p<0.001). Detailed statistics, exact p values and sample sizes for all comparisons are provided in **Supplementary Table 1.**

To characterize the spatial extent of these dendritic domains, we determined the physical boundaries of each turnover zone by identifying the contiguous cluster of turnover events flanking each density peak. Starting from the peak center, boundaries were defined by the outermost events connected by inter-event distances of <5μm (**Fig. 3f**). Turnover zone widths ranged mostly from 2.5 to 16 µm, with a mean of 6.5 µm (**Fig. 3g**), closely matching the ∼5-10-µm neighborhoods over which clustered synaptic plasticity has been reported^4,5,11,15,53^. Importantly, however, the zones reported here were defined based on spine turnover events and not the proximity of neighboring spines (i.e., clustering). Mapping turnover zones across all imaged segments illustrates that spine turnover was concentrated within a small number of such domains per dendrite, each hosting a range of gain and elimination events (**Fig. 3h**).

Because adaptation unfolds over many sessions, we next asked whether these zones are transient or persistent. We compared turnover zones detected from events in the early phase (sessions 0 to 4) with those detected from all events across the adaptation period (sessions 0 to last). The majority (59.3%) of zones present across the full period were already detectable in the early phase, while a minority (40.7%) emerged only once later events were included **(Fig. 3i**). The location of persistent turnover zones was highly stable, with their centers shifting on average ∼2 µm (**Fig. 3j**). Because the full-period analysis includes early-phase events, persistence is in part expected. However, the minimal displacement of zone centers indicates that subsequent turnover did not relocate the early zones. Zones emerging later occupied significantly narrower spans than persistent zones (median: 2.44 µm vs. 6.80 µm; mean: 3.79 µm vs. 8.35 µm; Mann–Whitney U test, p = 0.0146; **Fig. 3k**), consistent with localized remodeling initiating within tight dendritic windows. Alternatively, this difference may reflect the shorter period over which events contributing to these zones could accumulate. Turnover zones therefore constitute spatially stable domains that are largely established early after the switch and are re-used throughout adaptation.

These properties distinguish turnover zones from the turnover “hotspots” described in retrosplenial cortex by Frank et al. (2018)^11^. In that study, hotspots were dendritic segments with elevated spine turnover before learning, which predicted where learning-related clustered spine addition would subsequently occur. Turnover zones, by contrast, are resolved at sub-segment scale (mean width 6.5 µm, with up to two zones per segment), are defined from both spine gains and losses occurring during learning, and remain spatially stable across sessions, with most zones established shortly after the switch and re-used throughout adaptation. Rather than reflecting a segment-wide propensity for clustered spine formation, turnover zones thus represent a persistent, fine-scale organization of remodeling within individual dendrites.

Given the spatially restricted nature of turnover zones, we finally asked how formation and elimination events are distributed relative to these zones across the learning sessions following the switch. Because turnover zones were defined from the spatial density of these events, their concentration within zones is partly expected by construction. Evidence that turnover is concentrated beyond chance is provided by the permutation analyses in which zones were detected on shuffled data (**Fig. 3c, d**). We therefore used the present analysis to examine the composition of within-zone remodeling across event types and phases. Both formation and elimination events were enriched within turnover zones in the early and late phases, with >50% of events occurring within zones, compared with ∼17% expected if events were distributed randomly across valid spine locations (**Fig. 3l**). The proportion of elimination events occurring within zones was higher in the late than in the early phase (77.6% vs. 63.5%), although this comparison may partly reflect differences between phases in the number and extent of zones and in the events included in each analysis. These results indicate that gains and losses share the same restricted dendritic domains, and suggest that eliminations become increasingly confined to these domains as adaptation proceeds.

### Clustered formation and clustered elimination of spines within dendritic turnover zones

Having localized spine remodeling to stable dendritic turnover zones, we asked what is built and removed within these domains during flexible learning. In particular, whether newly formed spines are organized into spatial clusters, whether their organization is retained, and whether elimination is targeted to the same dendritic domain. Defining a cluster as two (or more) newly formed spines arising within 5 µm of one another^11,13^, we found that new spines frequently arose adjacent to pre-existing spines, forming extended groups of 3-8 spines. As expected, these groups showed a higher local spine density than randomly positioned windows on the same dendrite (**Fig. 4a**; Switch, 24 clusters, p=0.0025). We therefore tested clustering against chance separately using a permutation approach (**Fig. 4c**).

**Fig. 4.**
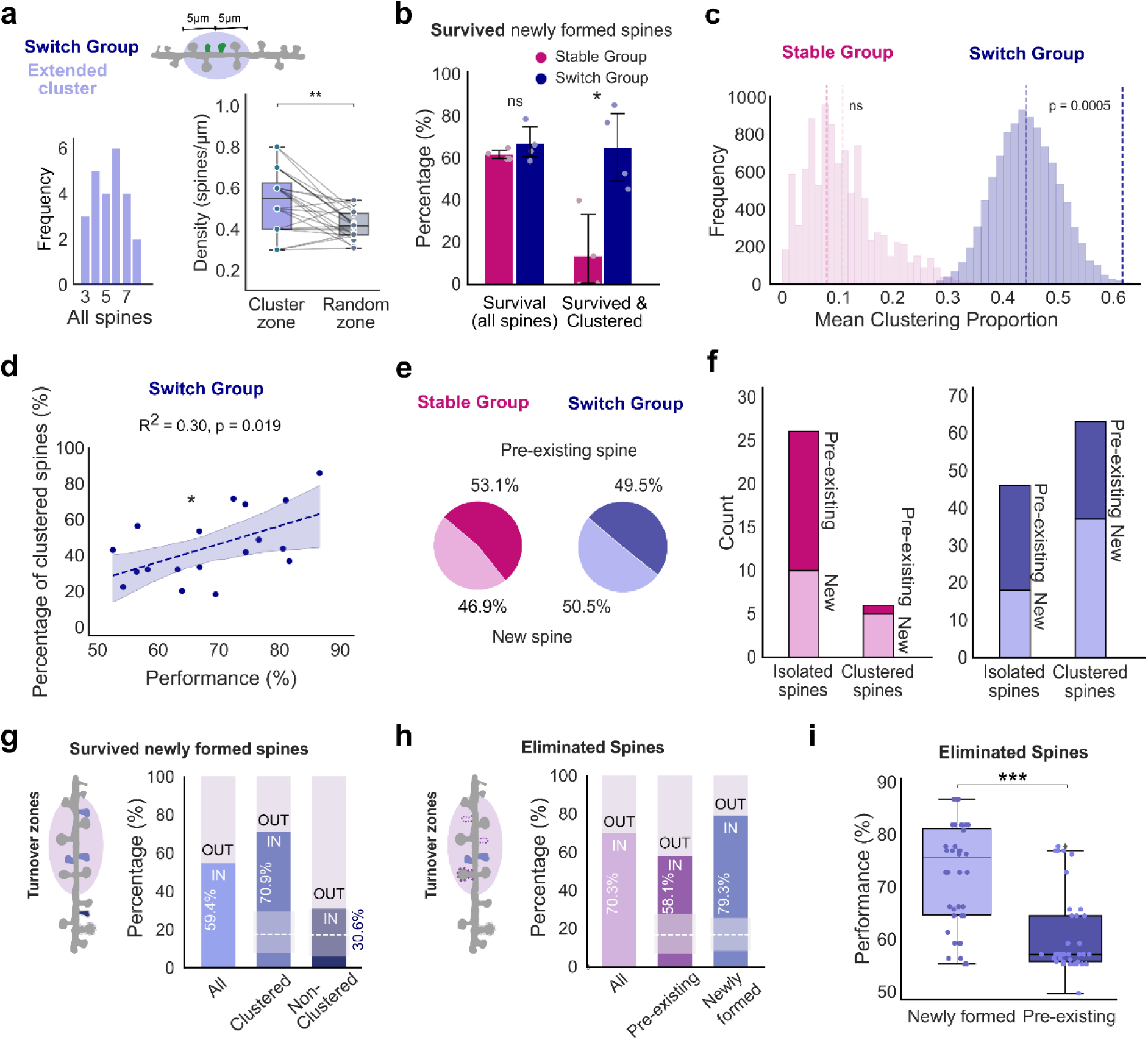
Clustered spine formation and elimination are coupled during flexible learning. **a**, Total spine density in cluster zones compared to a spatial null model. Total spine density inside a 5μm radius window centered on the cluster peak) was significantly higher than the expected density across 500 randomly positioned windows of identical width along the same dendritic branch (Switch, n = 24 clusters; p = 0.0025, two-tailed paired Wilcoxon signed-rank test). **b,** Survival of all newly formed spines (percentage persisting to the last imaging session) in the Stable and Switch groups; survival did not differ between groups (t=−0.84, p=0.43, t-test). Proportion of surviving newly formed spines belonging to a cluster (within 5 µm of another surviving new spine); the clustered fraction was significantly greater in the Switch group (t=−3.09, p=0.023, t-test). **c,** Observed clustered fraction of surviving new spines versus a null distribution generated by shuffling new-spine positions across biologically valid locations (10,000 permutations, density conserved). Clustering exceeded chance only in the Switch group (Switch, observed 0.62 vs. null 0.44, p=0.0005; Stable, observed 0.08 vs. null 0.10, p=0.97; Mann–Whitney U test). **d,** Proportion of clustered new spines as a function of behavioral accuracy across the adaptation period in the Switch group (linear regression, R^2^=0.30, p=0.019). **e**, Proportion of eliminated spines that were pre-existing versus newly formed, in the Stable (pre-existing 53.1%, new 46.9%) and Switch (pre-existing 49.5%, new 50.5%) groups; both populations were lost in roughly equal proportions. **f**, Spatial configuration of eliminated spines: Switch-group losses were predominantly clustered, whereas Stable-group losses were predominantly isolated. **g**, Localization of surviving newly formed spines relative to turnover zones, by population; Survived and clustered newly formed spines fell within turnover zones (70.9% inside), whereas the surviving, non-clustered new spines were non significantly enriched relative to turnover zones (30.6% inside). Dashed lines indicate chance levels (Monte Carlo permutations) for both groups: mean values are ∼17% for both groups with range of: [7-27%] for survived and clustered newly formed spines and [6-28%] for survived and non-clustered newly formed spines (paired Wilcoxon signed-rank test, survived and clustered: p<0.0001, survived and non-clustered: n.s.). **h**, Localization of eliminated spines relative to turnover zones. Newly formed eliminated spines as well as pre-existing eliminated spines fell predominantly within turnover zones (79.3% and 58.1%, respectively). Dashed lines indicate chance levels (Monte Carlo permutations) for both groups: mean values are ∼17% for all groups with range: [10-25%] eliminated pre-existing spines and [9-26%] for eliminated newly formed spines (paired Wilcoxon signed-rank test, both groups: p<0.0001). **i**, Behavioral performance at the time of elimination for pre-existing versus newly formed spines in the Switch group. Pre-existing spines were eliminated predominantly when performance was low (early post-switch), whereas newly formed spines were eliminated across the full performance range (p<0.0001). Box plots show median and interquartile range; shaded regions and error bars indicate ±1 SD. Detailed statistics, exact p values and sample sizes for all comparisons are provided in **Supplementary Table 1**.

The maintenance of these clusters, however, depended on behavioral demand. The overall survival of newly formed spines (at the end of the adaptation period) did not differ between groups (**Fig. 4b**; t=−0.84, p=0.43), but the proportion of surviving new spines that belonged to a cluster was markedly higher in the Switch group (**Fig. 4b**; t=−3.09, p=0.023). Because the clustered fraction depends on the number of surviving new spines per segment, we compared it with a null model in which new-spine positions were shuffled across biologically valid locations while conserving their number. Surviving spines clustered above chance only in the Switch group (Switch, observed 0.62 vs. random 0.44, p=0.0005; Stable, observed 0.08 vs. random 0.10, p=0.97; **Fig. 4c**). The proportion of clustered new spines further scaled with behavioral accuracy across the adaptation period (**Fig. 4d**; R^2^=0.30, p=0.019), whereas in the Stable group, where performance remained high throughout the imaged sessions, no such scaling was observed (**Extended Data Figure S5**). Clustered new spines are therefore preferentially stabilized when the animal must acquire a competing rule, linking their maintenance to the demand for flexibility rather than to high performance per se.

Surviving clustered spines were also concentrated within the turnover zones identified above: 70.9% of clustered new spines that persisted to the end of adaptation lay within a turnover zone, compared with ∼17% expected if spines were distributed randomly across valid spine locations (one-sided Wilcoxon signed-rank test, p < 0.0001; **Fig. 4g**). Surviving new spines that did not belong to a cluster showed no corresponding enrichment (30.6% inside vs. 17% chance with range [6-28%]; n.s.; **Fig. 4g**), indicating that these zones are selectively hosting those spines that (emerge and) survive in clusters rather than the isolated surviving population.

We next asked how spine elimination was organized relative to this formation, and specifically whether the domains hosting newly formed spines also hosted loss of spines. Pre-existing and newly formed spines were eliminated in roughly equal proportions in both groups (**Fig. 4e**), indicating that the switch drove bidirectional remodeling of the synaptic population rather than a selective gain or loss of one class of input. In the Switch group, eliminated spines were predominantly clustered, whereas in the Stable group they were predominantly isolated (**Fig. 4f**), indicating that elimination, like formation, is spatially organized under competitive demand.

Both eliminated populations were preferentially located within turnover zones, though to markedly different degrees. Newly formed eliminated spines fell predominantly within these zones (79.3% inside vs. 17% chance; paired Wilcoxon signed-rank test, p<0.0001; **Fig. 4h**) and pre-existing eliminated spines were also significantly enriched (58.1% inside vs. 17% chance, range [7-28%]; p<0.001; **Fig. 4h**). As for surviving spines (**Fig. 4g**), this enrichment is in part expected by construction, because elimination events contributed to zone detection. The effect is likely strongest for newly formed eliminated spines, each of which contributes both a formation and an elimination event at the same location, and this may account for part of their higher within-zone fraction. Pre-existing eliminated spines, by contrast, each contribute a single event, as do non-clustered surviving new spines, which were not significantly enriched (**Fig. 4g**). Their enrichment is therefore less readily explained by the zone-detection procedure alone, although clustered losses (**Fig. 4f**) may contribute multiple nearby events. Taken together, these results suggest that the same zones host the elimination of pre-existing spines, which may have supported the prior rule, alongside the formation, clustering, and elimination of spines added during acquisition of the new rule. Old and new spines would thus occupy a shared dendritic territory rather than separate ones, consistent with a single dendritic compartment being reused across competing behavioral contingencies rather than distinct territory being recruited for each.

Spine loss was also temporally structured. Pre-existing spines, present before the switch and therefore candidates for encoding the prior rule, were eliminated mostly during the early post-switch sessions, when performance was lowest and the former strategy was still being suppressed. Newly formed spines, by contrast, turned over more uniformly throughout adaptation (**Fig. 4i**; p<0.0001), as reported for motor learning^5^. Together, these results indicate that turnover zones function as general loci of structural spine turnover during adaptation: they host the formation and elimination of transient new spines, and they account for a substantial share of pre-existing spine elimination. This co-localization, together with the early timing of pre-existing spine loss, is consistent with the re-use of dendritic substrate as the prior rule is replaced, a hypothesis we test using the network model described below.

### A biophysically constrained circuit model reveals the representational economy of dendritic reuse

Longitudinal imaging localizes where and when synaptic remodeling occurs, but it cannot reveal which of the eliminated spines had encoded the prior rule, nor can it test what the spatial reuse of a compartment contributes to circuit-level representation. To address these questions, we built a biologically constrained model of an M2 microcircuit (**Fig. 5a**). The model incorporates multicompartmental pyramidal (PYR) neurons and three inhibitory populations (PV, SST, VIP), receiving rule-specific feedforward (FF1, FF2) and feedback (FB1, FB2) inputs for the two modalities, with a reward signal that modulated the PYR apical compartment and VIP interneurons. As the model represents a higher order cortical circuit, feedforward inputs were assumed to carry rule-specific, not cue-specific, information. The latter are assumed to be encoded in primary cortical areas. Learning was governed by three mechanisms acting on distinct timescales: calcium- and plasticity-related-protein-dependent LTP/LTD with a learning rate that scaled inversely with synaptic strength, structural turnover (spine formation and elimination), and homeostatic scaling (**Fig. 5b**; Methods). Importantly, the reward signal does not explicitly enter the rules governing synapse formation or elimination. Formation is triggered by sustained high local dendritic Ca^2+^ and synaptic activation, and elimination follows from a loss of activation together with insufficient Ca^2+^ support.

**Fig. 5.**
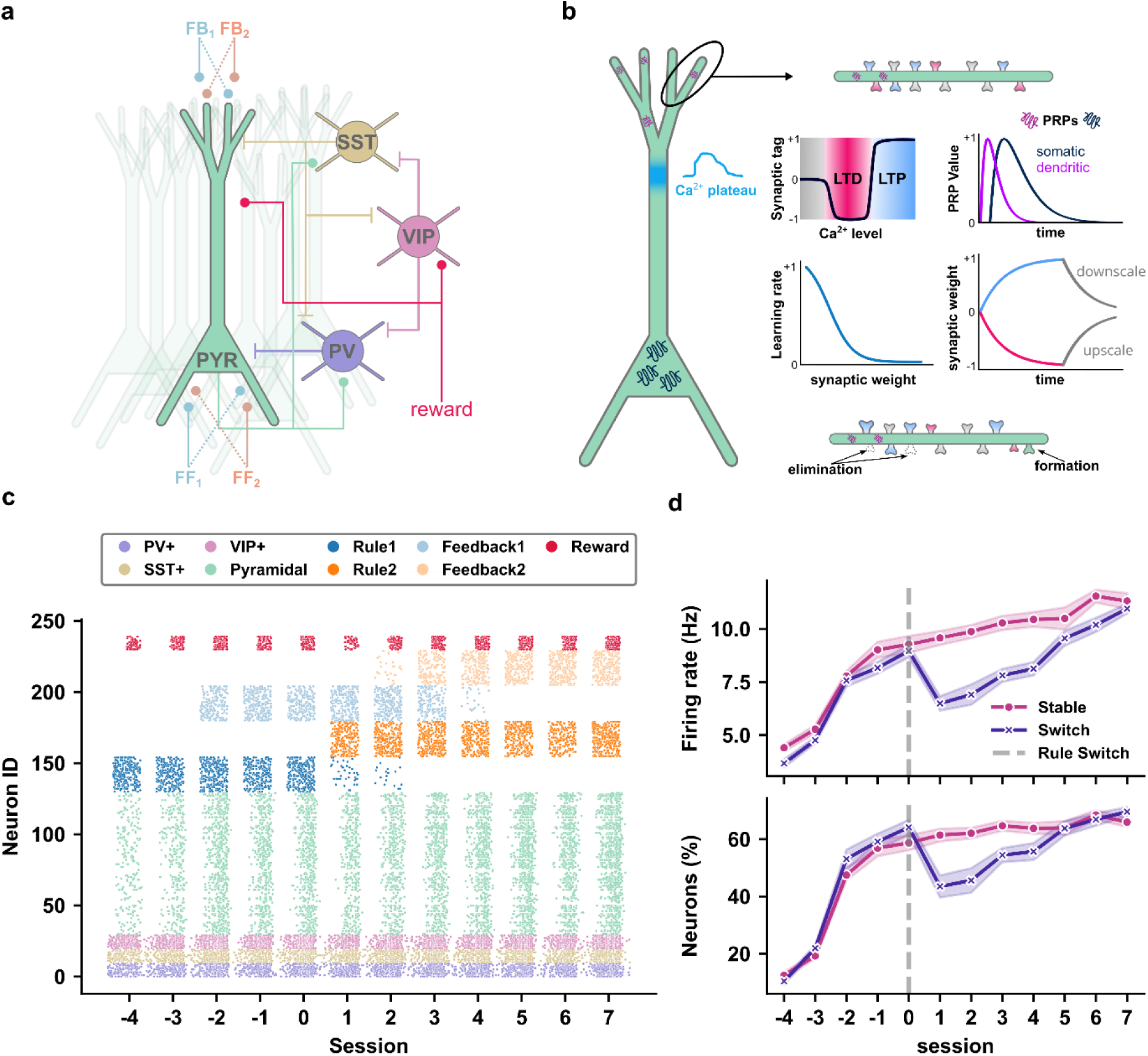
A biologically constrained M2 network model reproduces the population dynamics of cross-modal rule-switching. **a**, Schematic of the network microcircuit. Multicompartmental excitatory pyramidal (PYR) neurons interact with local PV, SST and VIP interneurons. The network receives rule-specific feedforward (FF1, FF2) and feedback (FB1, FB2) inputs; the reward signal modulates the PYR apical compartment and VIP interneurons. **b**, Learning rules. Functional plasticity follows a BCM-like synaptic tag-and-capture rule in which a sliding, calcium-dependent threshold separates LTD from LTP; the model further incorporates the spatiotemporal dynamics of plasticity-related proteins (PRPs) across somatic and dendritic compartments (modelled as alpha functions), a synapse-specific learning rate inversely proportional to synaptic weight, and homeostatic synaptic scaling (upscaling of depressed and downscaling of potentiated synapses). Structural turnover adds and removes spines according to local activity. **c**, Representative raster of network activity across learning sessions (Sessions −4 to 7) for the Switch condition; colors denote cell types and input streams (legend). **d**, Global network dynamics for the Stable (magenta) and Switch (purple) conditions across sessions. Top: mean population firing rate (Hz). Bottom: percentage of active neurons (firing rate > μ + 1σ, with μ and σ computed from the first two sessions as baseline). Both measures rise across acquisition and, in the Switch condition, show a transient depression immediately after the rule switch (dashed vertical line, Session 0) before recovering, paralleling the experimental trajectory (cf. Fig. 2b). Shaded regions indicate ±1 SEM across independent network simulations (20 seeds per condition).

We calibrated the model by adjusting its parameters until the activity of the model paralleled the behavioral performance of rodents and the average rates of spine gain, loss and turnover it produced matched those measured *in vivo* (**Fig. 6a**; cf. **Fig. 2e**). With these values fixed, no further parameters were tuned.

**Fig. 6.**
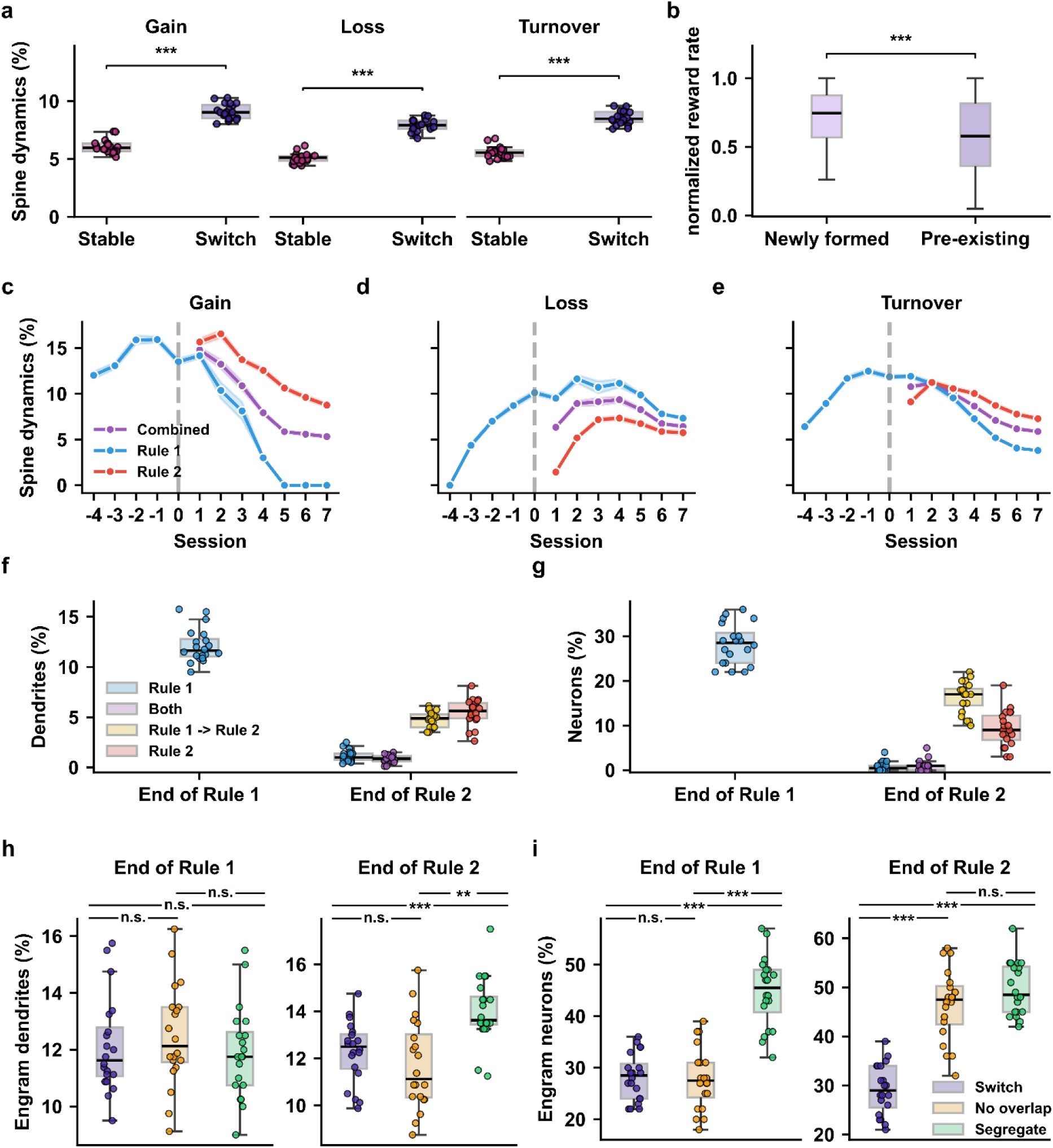
The model resolves the rule identity of eliminated synapses and reveals the representational economy of dendritic reuse. **a**, Calibration of the model. Plasticity parameters were adjusted to match in vivo turnover magnitudes of spine gain, loss and turnover (% of total spines) produced by the Switch and Stable networks (cf. Fig. 2e); with these values fixed, no further parameters were tuned. The Switch network shows significantly elevated dynamics across all three metrics. **b**, Normalized reward-input firing rate during the lifetime of newly formed and pre-existing spines in the Switch network, used as the model’s correlate of elimination timing. Newly formed spines were active during higher-reward periods than pre-existing spines at the time of their removal, reproducing the in vivo bipartite elimination pattern (cf. Fig. 4i). **c–e**, Pathway-resolved trajectories of spine gain (**c**), loss (**d**) and turnover (**e**), expressed as a percentage of total spines across sessions, shown separately for Rule 1 (blue), Rule 2 (red) and their sum (Both, purple). The dashed line at Session 0 marks the switch; Rule 2 and Both dynamics emerge only post-switch. Rule 1 synapses are preferentially eliminated early after the switch, coinciding with the rise in Rule 2 gain, while the combined trajectory remains relatively stable. Solid lines, mean; shaded regions, ±1 SEM across simulations. **f**, Percentage of dendrites classified as Rule 1-selective (blue), Both (purple), transitioning Rule 1 → Rule 2 (yellow) or Rule 2-selective (red) by combined structural and functional criteria (Methods), at the End of Rule 1 (left) and End of Rule 2 (right). Note the reduction in Rule 1 encoding dendrites (blue and purple) at the end of Rule 2. **g**, Percentage of neurons classified into the same four categories as in (f). Note the elimination of Rule 1 encoding neurons (blue and purple) at the end of Rule 2. **h**, Dendritic engram size, namely the percentage of dendrites classified as rule-specific or mixed-selective, in the intact Switch network (purple), the No-overlap condition (yellow; mixed-selective dendrites prevented) and the Segregate condition (green; Rule 1 and Rule 2 synapses forced onto non-overlapping compartments), at End of Rule 1 (left) and End of Rule 2 (right). By the end of Rule 2 the No-overlap network is indistinguishable from the intact network, and only full segregation expands the dendritic engram. **i,** as in **(h)**, for neuronal engram size, namely the percentage of neurons recruited as rule-specific or mixed-selective. Both constraints enlarge the neuronal engram by the end of Rule 2. Box plots show median and interquartile range; whiskers extend to minimum and maximum values. Statistical comparisons: Bonferroni-corrected Mann–Whitney U tests (**a**), Mann–Whitney U test (**b**), Kruskal–Wallis followed by Bonferroni-corrected pairwise Mann–Whitney U tests (**h**, **i**). *p<0.05, **p<0.01, ***p<0.001, n.s., not significant. Detailed statistics, exact p values and sample sizes for all comparisons are provided in **Supplementary Table 1**.

Driven by a reward signal that followed the measured learning curve and the respective FF and FB inputs for each rule (as shown in **Fig. 5c**), the model produced population-level signatures of the task. Both the mean pyramidal firing rate and the fraction of active neurons rose across acquisition and, in the Switch condition, showed a transient reduction immediately after the rule switch before recovering, aligning with the behavioral experimental trajectory (**Fig. 5c, d**; cf. **Fig. 2b**). The model thus recapitulates, at the population level, animal behavioral trajectories, providing an *in silico* testbed with which to dissect the subcellular reorganization that imaging cannot resolve.

### Dendritic reuse tracks the identity of eliminated synapses and confers representational economy

We first asked whether the model captured the temporal asymmetry of elimination observed *in vivo*: newly formed spines turned over throughout adaptation, whereas pre-existing spines were eliminated most heavily during the early post-switch period. In the model, the switch was implemented as a change in rule-specific input: Rule 2 feedforward inputs were introduced, while Rule 1 inputs continued at reduced firing rates for 3 sessions. This reduction is an imposed assumption representing attentional down-weighting of the sound cues, which remained present but irrelevant after the switch^54,55^. Deprived of this drive, and of the reward-driven apical depolarization that would otherwise sustain them, established Rule 1 synapses fell below the threshold for structural maintenance and were eliminated during the early sessions. Newly formed spines, by contrast, were still being actively driven by the incoming rule. Consequently, at the time of their removal, newly formed spines had been active during higher-reward periods than pre-existing spines (**Fig. 6b**), reproducing the *in vivo* pattern (**Fig. 4i**). As both the reduction of Rule 1 input and the reward signal were imposed, this asymmetry reflects how the plasticity rules respond to the switch protocol, rather than an independent prediction of its timing.

Because every synapse in the model carries its pathway label, the model can also address a question that *in vivo* imaging cannot: which rule the eliminated synapses had served. We therefore tracked formation and elimination separately for synapses from Rule 1 and Rule 2 input pathways. We found that Rule 1 synapses were eliminated mainly during the early post-switch period, coinciding with the rise in Rule 2 formation, whereas Rule 2 synapses turned over uniformly thereafter. Combined turnover remained relatively stable across the switch (**Fig. 6c-e**). Because Rule 1 input was reduced after the switch, the early loss of Rule 1 synapses is in part a consequence of the imposed protocol. Its temporal coupling with Rule 2 formation, however, indicates that within the calibrated network the removal of prior-rule synapses proceeds in step with the construction of the new representation. This provides a biophysically plausible account of the *in vivo* observation that pre-existing spines are lost early after the switch (**Fig. 4i**), consistent with, but not demonstrating, their encoding of the prior rule.

We next examined how these synaptic changes reorganized dendritic and neuronal selectivity. Classifying dendrites and neurons by combined structural and functional criteria (see Methods) as Rule 1-, Rule 2-, mixed- (Both), or transitioning-selective, we found that at the end of Rule 1 both populations were predominantly Rule 1-selective (**Fig. 6f, g**). The rule switch reorganized this pattern: Rule 2-selective dendrites emerged alongside a population of mixed-selective dendrites (**Fig. 6f**), underpinned by class-specific structural dynamics that exceeded the unclassified baseline (**Extended Data Fig. S6**). Mixed-selective populations arose at proportions above those expected from independent allocation of the two rules, at both the dendritic and the neuronal level (one-sided Wilcoxon signed-rank test; dendrites: p < 0.0001; neurons: p < 0.0001; **Extended Data Fig. S7**). This overlap is driven by the model’s plasticity rule, which adds synapses preferentially to compartments with sustained high Ca^2+^, such as those already strongly driven by Rule 1 inputs. Reuse therefore emerged from activity-dependent synapse formation, even though the architecture permitted the two rules to occupy separate compartments.

Because neuronal classification is derived from dendritic allocation, the above-chance overlap at the neuronal level is at least partly inherited from the dendritic level. By the end of Rule 2, Rule 1-selective neurons were almost entirely lost (**Fig. 6g**), whereas Rule 1-selective dendrites, although reduced, persisted (**Fig. 6f**). The persistence of a structural Rule 1 trace at the dendritic level could provide a substrate for the rapid relearning observed when animals return to the original rule ^6,56^.

Finally, we tested whether dendritic compartment reuse minimizes the dendritic and/or the cellular engram size across the successive rules. Under an identical reward schedule, we compared the intact Switch network against two perturbation models that prevented reuse: a “No-overlap” condition that blocked Rule 2 formation on dendritic compartments already classified as Rule 1 specific (preventing mixed-selective dendrites), and a “Segregate” condition that forced Rule 1 and Rule 2 synapses onto strictly non-overlapping compartments. Both constraints acted as intended, abolishing mixed-selective and transitioning dendrites by the end of Rule 2 (**Extended Data Fig. S8a**). Mixed selectivity nonetheless re-emerged at the neuronal level, where neurons combined the two rules across separate dendrites (**Extended Data Fig. S8b**). Neither constraint altered the population firing rate or the fraction of active neurons (**Extended Data Fig. S9**), indicating that overall network activity was preserved.

The constraints instead affected how rule-related synapses were distributed across dendrites and neurons, and the two levels dissociated. At the end of Rule 1, dendritic engram size did not differ between networks (**Fig. 6h**; Kruskal–Wallis test, p = 0.49). By the end of Rule 2, the No-overlap network still did not differ from the intact network (p = 0.84), whereas the Segregate network recruited more dendrites than either (vs. intact, p < 0.001; vs. No-overlap, p = 0.002; Kruskal–Wallis test, p < 0.001, followed by Bonferroni-corrected pairwise Mann–Whitney U tests). At the neuronal level, the No-overlap network matched the intact network at the end of Rule 1 (p = 1.0) but recruited significantly more neurons by the end of Rule 2 (p < 0.001; **Fig. 6i**). Therefore, preventing mixed-selective dendrites left the number of recruited compartments unchanged while distributing them over substantially more neurons. Because the fraction of active neurons was unaffected (**Extended Data Fig. S9**), this expansion reflects the spread of rule-specific dendrites across a larger set of neurons rather than the recruitment of additional active cells. The Segregate network, on the other hand, already showed a larger engram at the end of Rule 1 (**Fig. 6h, i**), as expected from partitioning the available compartments before learning. The No-overlap condition, by contrast, acted only after the switch: it did not differ from the intact network at the end of Rule 1 and diverged only during acquisition of Rule 2, isolating the effect of preventing reuse. Dendritic compartment reuse is thus not merely permissible but representationally economical, because enforcing spatial segregation between the two rules expands the dendritic and/or the neuronal population the network must recruit to hold both.

Together, the imaging and the model converge on a single account. The turnover zones that imaging localizes as stable, reused turnover domains correspond, in the network model, to compartments whose reuse across competing rules allows restructuring of the engram in response to behavioral demands, while conserving its overall size. This is an important representational saving that emerges from a model calibrated only to the magnitude of turnover, and that our spine imaging measurements could not have revealed.

## Discussion

Adaptation to a changed association raises a fundamental question: how are existing synaptic connections reorganized as new learning proceeds? By combining longitudinal two-photon imaging of M2 apical tuft spines with a biologically constrained, dendrite-aware network model, we show that spine formation and elimination are concentrated within persistent local dendritic domains during behavioral updating. In this account, spatially confined dendritic turnover zones act as reusable structural workspaces: they are domains in which new, rule-specific synapses are formed while obsolete ones are removed. The model suggests that sharing compartments across successive rules reduces the representational footprint. M2 inactivation impaired rule-switching, whereas no significant impairment was detected during rule maintenance or during acquisition of the light rule without competing cues (**Extended Data Fig. S2b, e**). Synaptic remodeling in M2 thus accompanies a form of behavioral updating that depends on this region.

Learning-induced plasticity concentrating within high turnover dendritic zones aligns with a substantial body of work showing that new spines form in spatial clusters and integrate with potentiated neighbors^4,5,13,15,18^. Whereas Frank et al.^11^ showed that segment-level turnover before learning predicts the location of subsequent clustered spine addition, we followed the sub-segment organization of spine formation and elimination during the updating of an established association. These remodeling domains spanned 2.5-16 µm (mean 6.5 µm), a scale comparable to short dendritic neighborhoods implicated in clustered synaptic plasticity. Together, these findings suggest that local dendritic domains can support both the establishment and subsequent reorganization of synaptic connections. These zones were also present above chance in the Stable group under matched sensory and motor demands, indicating that spatially concentrated turnover is not restricted to rule-switching. The switch therefore appears to increase the rate of remodeling these zones sustain, not their existence. Turnover zones were also spatially stable: most zones detected across the full adaptation period were already present in the early post-switch sessions, and their centers shifted by only ∼2 µm. Although this persistence is partly expected, because the full-period analysis includes early events, the minimal displacement suggests that local dendritic domains provide a stable spatial framework within which individual synaptic connections are replaced during adaptation.

The same zones that hosted new-spine formation and clustering were also enriched for the elimination of pre-existing spines, the population most likely to have supported the previous rule. Because turnover zones were defined from these same events, this co-localization is partly expected by construction; pre-existing eliminated spines, however, each contribute only a single event, yet were enriched well beyond chance, unlike surviving non-clustered new spines. These findings are supported by our modeling results. Pre-existing synapses eliminated after the switch were predominantly Rule 1 synapses, and their loss coincided with the formation of Rule 2 synapses on the same compartments. Together, the imaging and modeling support an account in which local dendritic domains accommodate a new association through both the addition of new connections and the removal of existing ones.

The timing of spine elimination further distinguished the remodeling of pre-existing and newly formed connections. Pre-existing spines were lost mainly early after the switch, when performance was lowest and the former strategy was presumably still being suppressed, whereas newly formed spines continued to turn over throughout adaptation. Part of this difference may reflect availability, since newly formed spines can only be eliminated after they appear. It is nevertheless consistent with an early phase in which established connections are removed, followed by continued formation and elimination as the new association is acquired. Local remodeling may therefore comprise overlapping processes with different time courses rather than a uniform increase in turnover.

The model allows us to address two mechanistic questions that our imaging experiments cannot answer. First, do the old and new rules engage the same dendritic compartments? Although the architecture allowed the two rules to occupy separate compartments, they recruited overlapping compartments more often than expected by chance, a consequence of the activity-dependent formation rule, which favors compartments already strongly driven by the first rule. This is consistent with the *in vivo* observation that the same turnover zones host both the loss of pre-existing spines and the formation of new ones, and with the interpretation that early-eliminated pre-existing spines encoded the prior rule. Second, what is the computational benefit of compartment reuse? When Rule 2 synapses were prevented from forming on compartments used by the first rule, the network acquired the second rule with a similar number of rule-selective dendrites. However, these dendrites were distributed over a larger population of neurons, without any change in overall activity. Within the model, reuse is therefore representationally economical at the neuronal level, consistent with the larger coding capacity conferred by mixed selectivity^33^. This is a distinct saving from the capacity benefit that elevated pre-learning turnover provides when memories accumulate serially^11^; here, the saving arises because successive rules share common dendritic territory and obsolete synapses are removed. Finally, whereas Rule 1-selective neurons were almost entirely lost by the end of Rule 2, a partial dendritic trace of Rule 1 persisted. This suggests a potential mechanism for the rapid relearning evidenced in animals upon returning to the original rule^6,56^.

One possible mechanism for the representational economy is local competition between neighboring synapses^57,58^. In our model, synapses on the same compartment share locally synthesized plasticity-related proteins, and homeostatic scaling conserves total synaptic weight, so that potentiation of new synapses weakens existing ones and brings them closer to elimination. As a result, new synapses can promote the formation of nearby spines^16^, while weak neighbors are pruned, plausibly through shared local resources^59^. Signals from areas such as orbitofrontal cortex^60^ could regulate these local interactions as reward expectations change. Whether such signals control the spatial pattern of spine formation and elimination remains to be tested.

Taken together with previous studies, our results suggest that the relationship between learned associations may influence whether they recruit shared or separate dendritic territory. Independent skills with non-overlapping demands likely recruit different dendritic segments that minimize interference^10^; cooperative memories formed close in time co-allocate to shared segments to bind recall^13,27^. Our findings suggest that updating competing rules can involve shared dendritic domains in which new connections form and pre-existing connections are removed. This raises the possibility that the balance between synapse formation and elimination helps determine whether shared dendritic territory links memories or supports replacing an established association with a new one.

Our findings extend the question of how memories share or separate neuronal ensembles^29–31^, to the remodeling of connections within dendrites, complementing evidence for engram-specific spine clustering in motor circuits^22^. Within this framework, the functional reorganization of M2 tuft compartments and the representational drift observed upon relearning the original rule^32^ could be the activity-level signature of the structural reuse we find here. Thus, structural and functional synaptic signatures observed in the two studies may reflect a common determinant of adaptive learning. Moreover, because this functional reorganization did not depend on local inhibitory control^32^, rule-dependent adaptation may be expressed at the level of pyramidal tuft dendrites and their synapses rather than inherited from a reconfiguration of local inhibition.

Several limitations should guide the interpretation of these conclusions. First, we did not measure the functional identity of eliminated pre-existing spines directly; the model predicts that early-eliminated spines preferentially encode the prior rule, but testing this prediction would require tracking both the structure and rule-related activity of individual spines. Second, turnover zones were identified from the same events whose localization we analyzed, so within-zone enrichment is partly expected by construction. The permutation analyses establish that turnover is concentrated beyond chance, but cross-validated designs, in which zones are defined from one phase or event type and tested on another, would provide a stronger test. Third, imaging was restricted to the second rule phase, leaving any overlap with turnover during initial acquisition unexamined. Fourth, M2 inactivation shows the region is required for rule-switching but does not show that turnover dendritic zones are necessary for behavioral updating. Fifth, the model is deliberately simplified to keep it tractable while capturing essential features: coarser compartments, session-wide plasticity updates, an imposed rather than earned reward and input schedule. Model outcomes concerning the timing and rule identity of eliminated synapses, therefore depend in part on this protocol. Finally, the model shows that dendritic reuse can reduce neuronal recruitment under these conditions; it does not establish whether reuse improves behavioral performance, learning speed or relearning.

Overall, our results support a view in which local dendritic domains a few micrometers long (mean 6.5 µm) provide a persistent framework within which synaptic connections are reorganized as learned associations change. The model identifies a potential computational benefit of this organization: reusing dendritic domains allows a new rule to be acquired with fewer neurons than when its synapses are kept separate from those of the previous rule. Reuse, however, comes at a price: in the model, the neuronal representation of the old rule was largely overwritten, the trade-off that defines catastrophic forgetting in artificial systems^61^. The most effective remedies, biological and algorithmic alike, work by protecting earlier-task weights while permitting change elsewhere^1^, often through overlapping-but-distinct dendritic subnetworks^62,63^. The partial dendritic trace of the old rule in our model may represent one form of such protection. Together, these results suggest that the capacity to update learned associations depends not only on which synapses change, but on how their formation and elimination are organized within the dendritic tree.

## Acknowledgments

The authors thank Prof. M Froudarakis, Prof. K Sidiropoulou, Prof. S Smyrnakis, Prof. L. Palmer, Prof. M Seghal, Dr. Naoya Takahashi and the members of the Poirazi lab for helpful discussions supporting this project. We thank Prof. T Komiyama and Dr. WJ Wright for sharing a GUI for extracting spine positions; A. Antoniadou and I-R Tzonevrakis for their contributions in the development of the computational model; Dr. S Spanou, M Protopapa, P Vouvopoulos and Prof. D Bazopoulou for their contribution in confocal microscopy. We thank Dr. R Makarov for his contribution to schematics used in this paper, and the Research Workshop at the Charité – Universitätsmedizin, Berlin for developing and manufacturing devices in the behavioral apparatus.

## Author contributions

Conceptualization for experimental work: I.P., A.P., P.P., M.E. L.; Methodology for experimental work: I.P., H.O., M.A.N., A.P., P.P., M.E.L.; Conceptualization and methodology for computational modeling: S.C., P.P., I.P.; Performed all experiments and analysis: I.P., S.C.; Funding acquisition: A.P., P.P.; Supervision: A.P., P.P.; Writing – original draft: I.P., P.P., S.C.; Writing – review & editing: all authors.

## Methods

### Animals

All procedures were approved by the Veterinary Directorate of the Prefecture of Heraklion (Crete) and conducted in accordance with the guidelines of the Greek Government and the FORTH Ethics Committee. C57BL/6 and Thy1-GFP-M mice (Tg(Thy1-EGFP)MJrs; The Jackson Laboratory, stock #007788), 3–6 months old, were housed at the IMBB-FORTH Animal Facility on a reversed 12 h light/12 h dark cycle (lights off 07:00–19:00), with all behavioral testing performed during the dark phase. Food and water were available ad libitum until the onset of water restriction (see Water restriction). Mice were randomly allocated to experimental groups, and no animals were excluded from analysis. C57BL/6 mice were used for the pharmacological inactivation experiments and Thy1-GFP-M mice for the longitudinal spine-imaging experiments.

### Surgery

Adult mice (∼P100) were anaesthetized with ketamine (100 mg/kg)/xylazine (10 mg/kg) and maintained on a thermal blanket in a stereotaxic frame (Stoelting). A lightweight head-post was fixed to the skull with RelyX and dental acrylic, leaving space for a cranial window. A 3-mm-diameter craniotomy was made over the secondary motor cortex (M2; AP +1.8 mm, ML +0.75 mm), the dura was left intact, and a 3-mm (No. 0) glass coverslip was placed and secured with cyanoacrylate and dental acrylic. For the muscimol experiments, two small openings were made in the coverslip over M2 bilaterally (AP +1.8 mm, ML ±0.75 mm) to permit repeated intracortical access across days using an Ultra Precise Digital Mouse Stereotaxic Instrument (Stoelting). The coverslip was then covered with silicone sealant (Kwik-Cast, World Precision Instruments) secured with glue. Carprofen (5 mg/kg, s.c.) was administered before the end of surgery and again two days later to manage postoperative pain.

### In vivo pharmacology

Before every post-switch training session, C57BL/6 mice were lightly anaesthetized with isoflurane (1.5–2%, constant flow 1 L/min). Through the two coverslip openings (AP +1.8 mm, ML ±0.75 mm), muscimol (1 µg/µl) or physiological saline vehicle (0.9%) was infused bilaterally at two depths (700 and 350 µm below the pial surface; 100 nl each, 200 nl per hemisphere) at 50 nl/min, using glass pipettes (Drummond, Cat# 5-000-2005) pulled and broken to a ∼10– 15 µm inner-diameter tip to minimize dural damage. Mice were randomized to receive saline or muscimol. The coverslip was resealed with silicone sealant and mice recovered in their home cage for 30 min before the behavioral task. After the final pharmacology session, fluorescent muscimol (BODIPY TMR-X conjugate, Thermo Fisher Scientific) was infused at the same sites and depths (100 nl each), and mice were perfused 30 min later to verify the spread of the infusion.

### Behavioral Setup

The head-fixed behavioral apparatus was adapted from ^42,48^. A Plexiglas air table perforated with evenly spaced holes directed compressed air upward to levitate a 15-cm-diameter PLA platform, allowing head-fixed mice to actively rotate the platform left or right, analogous to the treadmill or trackball used in other head-fixed setups. The platform carried three nose-poke ports (one central and two lateral choice ports), each with an LED, an infrared sensor/emitter, and a solenoid-controlled reward tube. Two speakers positioned ∼20 cm in front of the platform delivered auditory stimuli, and the LED within each choice port delivered visual stimuli. All hardware was controlled by Bpod (Sanworks), which read the nose-poke sensors, gated the reward solenoids, and presented the sensory stimuli; task parameters (e.g., the relevant rule and reward volume) could be adjusted online through its graphical interface.

### Water Restriction

Mice recovered from surgery for at least 4 days before water restriction began. For ∼5 days before behavioral testing, daily water intake was limited to 1–1.3 ml, delivered at a fixed time each day. Body weight was monitored daily; mice received supplementary water if their weight fell below 80% of baseline.

### Behavioral Training

Water-restricted mice were first trained on the sound rule (SR) while freely moving, to habituate them to the task and apparatus. Trials were self-initiated by a poke into the central port, which delivered a small water reward (0.55 µl), immediately followed by one of two auditory stimuli: an up-sweep (from 11 to 15 kHz) or a down-sweep (from 16 to 12 kHz). To facilitate acquisition, sweeps were initially presented directionally. Mice had to choose the left port after an up-sweep or the right port after a down-sweep to obtain a larger reward (5.5 µl) at the choice port; incorrect choices triggered a high-pitched tone (∼2 kHz). After ∼3 days, mice were habituated to head-fixation and to moving the floating platform (initially with experimenter assistance) while continuing on the SR.

Once mice controlled the platform reliably under head-fixation, task parameters were tightened: incorrect choices incurred a 2-s timeout before a new trial could begin; the response window following central-port exit was reduced from 20 s to 10 s; and reward volumes were set to ∼0.55 µl (central port) and ∼5.5 µl (choice ports). Withdrawal from the central port before a 250-ms minimum sampling time aborted the trial with a 2-s timeout. Sessions comprised ∼220 trials and lasted 20–30 min, varying with response times, timeouts and accuracy. When mice exceeded 75% correct on the SR for two consecutive sessions, an irrelevant visual cue (a light at the left or right port) was introduced as a distractor for one session; after mice maintained >75% correct with the distractor present, the contingencies were switched to the light rule (LR), under which the illuminated port predicted reward and the auditory stimuli became irrelevant.

### Behavioral performance and error classification

Session performance was defined as the proportion of correct trials among all completed trials. Within-session learning dynamics were characterized with an 80-trial sliding-window average advanced one trial at a time; for the first 80 trials, baseline performance was taken as the mean accuracy of trials 1-80.

To quantify adherence to each sensory modality independently of the active rule, we computed the probability of following each cue:

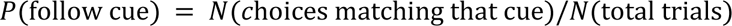

A choice was counted as “following the sound cue” if the mouse selected the side that the SR would have rewarded (e.g., the left port after an up-sweep), independent of the visual cue and of whether the trial was rewarded under the current rule; the probability of following the light cue was defined analogously. This metric allowed residual adherence to the now-irrelevant sound cues to be tracked after the switch.

Following the rule switch, incorrect choices were categorized based on their relationship to the new visual rule (Light Rule, LR) and previously learned auditory rule (Sound Rule, SR). On incongruent trials (conflicting visual and auditory cues), errors directed toward the port indicated by the sound cue were classified as perseverative errors, reflecting a perseverative choice to the former strategy. On congruent trials (aligned visual and auditory cues), errors were classified as random errors, consistent with neither the previous nor the current rule. Error types were normalized as percentages of total incongruent trials and congruent trials, respectively. This normalization isolates strategy competition from general task trial generator algorithm, ensuring that changes in error rates are independent of individual differences in overall incongruent and congruent count trials.

### Behavioral temporal dynamics

Reaction time was defined as the interval from cue onset to central-port exit. Movement time was defined as the interval from central-port exit to choose-port entry, during which the mouse rotated the platform. Sampling time was their sum and corresponds to the interval over which the stimulus was available to the animal. Trial-by-trial sampling times across all sessions and mice were fitted with a first-order linear regression, and the per-trial means were used to derive a single representative trajectory.

### Two-Photon Microscopy

Imaging was performed on a two-photon microscope (Thorlabs MM201, upgraded with galvo-resonant scanning), with GFP excited at 920 nm (Coherent Axon 920 TPC). Stacks were acquired through a 16× 0.8-NA water-immersion objective (Nikon, 3-mm working distance) at 1024 x 1024 pixels using Thorlabs acquisition software, with emission detected by a Hamamatsu GaAsP photomultiplier tube. All sessions were imaged ∼2 h after the corresponding behavioral session under less than half the induction dose of the ketamine/xylazine mixture. Apical dendritic segments of layer 5 pyramidal neurons were imaged within 200 µm of the cortical surface (likely within layers I and II/III); segments oriented largely in the imaging x–y plane with minimal z-projection were selected, and z-stacks were acquired at 1-µm steps to span each segment. Because of the limited axial resolution of in vivo two-photon microscopy, only protrusions extending laterally within the x–y plane were included in the analysis ^64^.

### Longitudinal imaging and spine flagging

The same dendritic segments were relocated across days using a navigational map established during the first imaging session. Raw z-stacks were projected to two-dimensional .tiff images in ImageJ and analyzed with Activity Viewer (https://github.com/wjakewright/Activity_Viewer), an interactive Python interface developed by Dr. W. Wright (Prof. T. Komiyama laboratory) for manual ROI drawing of spines and their parent dendrites from two-photon data. For the first session, the software extracted spine identities, flags and positions along each selected segment; for subsequent sessions, the new z-stack of the same segment was loaded together with the previous session’s ROIs, flags and coordinates to maintain per-spine longitudinal tracking. At session 0 all spines were flagged “pre-existing”; in later sessions spines could additionally be flagged “newly formed” or “eliminated” according to their presence in earlier sessions, as defined below. Imaged distal tuft dendritic segments in Layer 1 were traced along their parent apical trunks back to their somata in Layer 5, confirming their identity as L5 pyramidal neurons and excluding L2/3 apical dendrites.

Ketamine administration can produce an acute elevation in spine formation within the first 24-72 hours following initial exposure that subsequently plateaus, with net spine density and turnover settle into equilibrium following repeated administration, with fractional changes (ΔDensity/Baseline) stabilizing near zero ^65^. In our experimental design, all animals underwent at least 2 imaging sessions during the SR training phase to verify GFP expression of L5 pyramidal neurons, window clarity and to select the dendritic compartments based on the inclusion criteria (see Data Inclusion Criteria). Furthermore, both groups received identical anesthesia exposure, and spine turnover in the Stable group remained within typical physiological baseline ranges throughout the experiment.^8,64^

### Classification of Dendritic Spines

Spines were identified and tracked along dendritic segments following established protocols^8^. Representative images were generated as mean-filtered z-projections in ImageJ. Only segments yielding high-quality stacks across all sessions were quantified, ensuring that inter-session tissue rotation did not affect spine identification. Spine dynamics were scored by comparing sequential images of the same segment: spines present at the first session were flagged pre-existing and considered stable if they persisted across all sessions; a spine was scored eliminated if present in a previous session but absent in the next, and newly formed if absent in previous session(s) but present subsequently.

Spine formation rate was the number of newly formed spines in session *i* divided by the total spine count in session *i*; elimination rate was the number of eliminated spines in session *i* divided by the total spine count in session *i* − 1; each total rate is the mean across sessions. Spine density for a segment was the spine count in session *i* divided by segment length; densities were normalized to each segment’s baseline and averaged per mouse within 2-day time bins (binning was used to smooth noise and accommodate uneven inter-session intervals). Survival refers to newly formed spines that remained present until the last imaging session.

To verify the spatial stability of tracked spines, we computed the inter-session jitter for each spine, defined as *jitter* = |position(*n*) − position(*n* + 1)|. Maximum jitter median is ∼1μm. Spines whose maximum *jitter* across sessions exceeded 3 µm were flagged for manual re-checking and corrected or excluded; the mean position across sessions was used in all position-dependent analyses.

### Identification and analysis of dendritic turnover zones

Turnover zones —dendritic domains of high spine-event density (gains and losses)— were identified with a two-step pipeline combining sliding-window spatial analysis with permutation testing (**Fig. 3**). The spatial distribution of spine events along each segment was quantified by counting events within 5-µm bins and averaging across window steps to yield a mean spatial-density curve of spine events. Turnover zones were defined as local maxima of this curve, detected with a minimum spine-count threshold of 90th percentile of the cumulative signal value distribution across all processed dendritic profiles, and a minimum inter-peak distance of 10 µm, ensuring that peaks reflected distinct biological clusters rather than stochastic fluctuations ^4^. Dendritic branches exhibiting at least one verified high-amplitude signal peak were classified as structural turnover zones. Both the spatial span and number of turnover zones were robust to the choice of binning window (3, 5, 7, and 10 µm) (**Extended figure S4 e, f**).

To test whether these high turnover dendritic zones arose by chance, we ran a Monte Carlo permutation analysis. For each segment containing at least one zone, 1,000 randomized datasets were generated in which the observed number of spine events (gain or loss) was preserved but their positions were randomly re-assigned to all imaged event locations along the segment, conserving spine density. Each randomized dataset was passed through the identical sliding-window and peak-detection pipeline to build a per-segment null distribution. Observed and null distributions were compared on the mean number of peaks per segment, peak amplitude, and, for segments with ≥2 zones, the inter-peak distance, using the Mann– Whitney U test (or an independent t-test where normality held by Shapiro–Wilk).

To verify that the observed formation and elimination events are enriched inside the zones during both the early and late phases, we executed a Monte Carlo spatial permutation analysis (1,000 permutations). For every dendritic branch, we calculate the total number of observed new or eliminated events for each phase and randomly reposition them across all available spine positions along each dendritic branch. For each permutation, we measure the percentage of the events that were positioned by chance inside the turnover zones to build a null-distribution. Observed value and null distribution for each group were compared using the one-sided Wilcoxon signed-rank test. We used similar analysis (Monte Carlo spatial permutation analysis (1,000 permutations) to calculate all chance levels in groups: clustered, non-clustered survived newly formed spines and pre-existing, newly-formed eliminated spines.

### Analysis of Spine Clustering

#### All newly formed spines

For each segment, we performed a pairwise distance analysis over all newly formed spines tagged across sessions (whether or not they were later eliminated). A spine was “clustered” if it arose within 5 µm of another new spine and “isolated” otherwise. We computed the mean clustering percentage per segment, then per mouse, for each group. Local spatial organization was further characterized by nearest-neighbor distance (NND), the minimum distance from each new spine to any other new spine on the same segment.

#### Surviving newly formed spines

The same pairwise analysis was applied to newly formed spines that survived until the end of learning —defined as ≥2 sessions above 75% on the LR for the Switch group, or the last imaging session for the Stable group. A surviving spine was “clustered” if within 5 µm of another surviving new spine. Approximately 70% of surviving newly formed spines remained for 2 or more imaging sessions.

#### Null model for clustering

To test significance, the positions of newly formed spines were shuffled across biologically valid spine locations (pre-existing, eliminated or newly formed), conserving per-dendrite spine density and segment length. The clustered fraction (<5 µm) was computed per segment, averaged per mouse, and then across mice to give one data point per observed data or shuffle; in the shuffle the previous analysis was repeated 10,000 times to build the null distribution.

#### Clustering versus performance

To relate clustering to behavior, we tracked, session by session, the clustering of surviving newly formed spines (a spine scored clustered if within 5 µm of another new spine present at that session), yielding a session-specific clustering profile aligned to performance. Significance was assessed by linear regression for the Switch group only; the Stable group’s performance remained above threshold and stable, so a clustering-versus-performance percentage was reported instead ^11^.

#### Hierarchical clustering

Spatial clusters were identified with DBSCAN applied to longitudinal spine positions using a 5-µm distance threshold. Cluster size was the number of newly formed spines within a cluster. We also counted all new and pre-existing spines within ±5 µm of each cluster center; local density inside the cluster was this count divided by the maximum pairwise distance among those spines, and density outside the cluster was the number of remaining new and pre-existing spines divided by the length of the remaining segment.

#### Elimination clustering

Spatially clustered eliminations were identified from pairwise distances among eliminated spines only: an eliminated spine was flagged clustered if within 5 µm of at least one other eliminated spine, and classified as clustered or isolated while distinguishing newly formed from pre-existing populations. For robustness in **Fig.4i**, elimination analyses were run on two datasets: the full set of imaged dendrites, and the subset imaged on the day of the rule switch (session 0). For a small number of dendrites, the initial imaging session acquired at behavioral Session 1; these segments were excluded from the restricted dataset to confirm that analyzed “pre-existing” spines were established before the switch rather than transiently gained on the switch day.

### Data inclusion criteria

Inclusion was determined by two-photon image quality across sessions: (1) low background noise; (2) dendrites well isolated from other fluorescent structures; and (3) healthy morphology—segments showing a marked reduction in spine density, dendritic “smoothing”, or “blebbing” were judged unhealthy and excluded. Mice whose imaging quality degraded substantially over the experiment were excluded; otherwise, all data were included and no statistical outliers were removed. Because repeated imaging risks photodamage, dendritic health was monitored throughout, and imaging sessions were selected to sample defined performance milestones (∼50–60%, ∼65%, ∼75%, and above threshold, >75%). We included filopodia in all structural tracking and turnover analyses. In our dataset, filopodia protrusions comprised 18.18% of the analyzed spine pool, with 12.61% persisting across consecutive imaging sessions. All filopodia were therefore tracked across sessions and included in analysis to ensure a complete accounting of spine reorganization following the rule switch. Spines present in a session, undetected in an intermediate session, and subsequently re-emerging in the next session at the exact same location on the dendritic shaft (<1μm) were classified as stable spines.

### Histology

To assess muscimol diffusion in M2, mice were deeply anaesthetized with ketamine/xylazine after the behavioral experiment and perfused with 4% paraformaldehyde. Brains were post-fixed for 24 h at 4 °C, washed in glycerol-buffered saline, sectioned at 20 µm, and mounted on slides. Images were acquired at x4 magnification on a confocal microscope. The M2 region was localized using the mouse brain atlas (Franklin and Paxinos).

### Computational model

#### Network composition and neuronal models

The simulated microcircuit consists of 130 neurons, comprising a primary excitatory population and three distinct inhibitory interneuron populations. The excitatory pool contains 100 multicompartmental pyramidal (PYR) neurons. The inhibitory circuitry is composed of 30 interneurons, divided equally into three functional classes: 10 Parvalbumin-expressing (PV), 10 Somatostatin-expressing (SST), and 10 Vasoactive Intestinal Peptide-expressing (VIP) interneurons. While the PYR neurons possess complex dendritic morphologies to support localized spatial integration (2 basal compartments, 3 apical trunk, and 6 apical tuft compartments), the PV, SST, and VIP interneurons are implemented as single-compartment point neurons. The interneurons are governed by the leaky conductance-based Adaptive Integrate-and-Fire (CAdIF) model utilized for the PYR somatic compartments.

The somatic compartment, governed by the leaky CAdIF model, captures essential spiking behaviors such as adaptation and bursting. Below is the set of ordinary differential equations that describe the change of the membrane voltage of a compartment *s*:

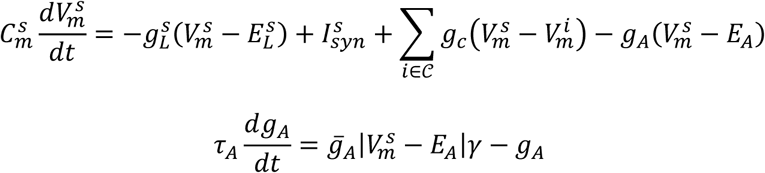

where *C*^*s*^_*m*_ denotes the capacitance, *g*^*s*^_*L*_ the leaky conductance, *E*^*s*^_*L*_ the resting potential, *g_A_* the adaptive conductance, *g̅*_*A*_ the maximal adaptation conductance, *E*_*A*_ the reversal potential of the adaptation, *γ* the steepness of the adaptation, *τ*_*A*_ the adaptation time constant, *g*_*c*_ is the coupling conductance between all neighboring compartments belong to neighborhood *C* (here denoted with superscript *i*), and *V*^*s*^_*m*_ the membrane voltage of the compartment. When *s* is any dendritic compartment, the adaptation parameter is set and kept at zero (i.e., no adaptation). When *s* is the somatic compartment, the spiking mechanism is described by:

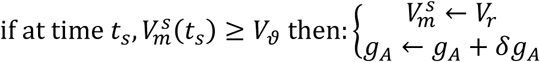

where *t*_*s*_ is a spike time, *V*_*ϑ*_ the spiking threshold, and *V*_*r*_ is the reset potential. *δg*_*A*_ is the spike-triggered adaptation, which describes the amount of increase in the adaptive conductance after each spike.

To replicate the non-linear processing of biological neurons, the dendritic tree is divided into distinct compartments implemented using the Dendrify framework^66^. These compartments are equipped with active mechanisms capable of generating local dendritic events, including sodium (Na^+^), calcium (Ca^2+^), and NMDA spikes. These mechanisms allow the model to perform complex spatial and temporal integration of inputs that go beyond the capabilities of standard point-neuron models. Furthermore, the model explicitly simulates intracellular calcium dynamics to bridge the gap between electrical activity and long-term structural changes. Each presynaptic spike triggers a transient calcium influx into the specific dendritic compartment where the synapse is located. This localized calcium signal acts as the primary driver for the plasticity rules, enabling the network to adjust its synaptic strengths based on the precise timing and location of input activity.

The calcium influx triggered by each presynaptic spike is modeled as a state-dependent process, governed by the following logistic (sigmoid) function:

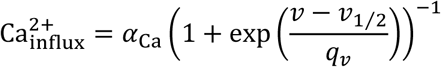

where *α*_Ca_ represents the maximum calcium influx (the upper asymptote of the curve), *v*_1⁄2_ is the half-activation potential (or midpoint), indicating the voltage *v* at which the influx reaches half of its maximum value, and *q*_*v*_ is the slope factor, which determines the voltage sensitivity or steepness of the sigmoid transition.

The model implements a multi-stage calcium signaling framework to capture the complex dynamics of intracellular Ca^2+^. This process is governed by three primary equations:

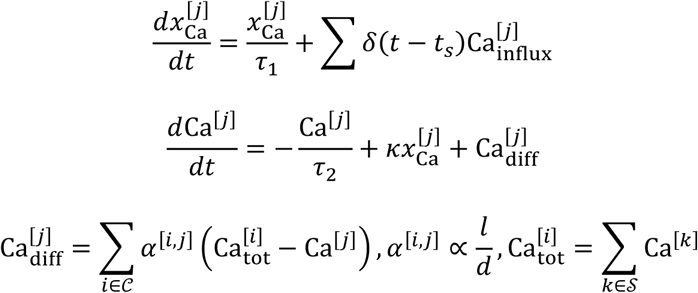

where *δ*(⋅) is the Kronecker delta function given by:

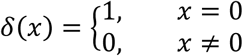

The temporal evolution of intracellular calcium is modeled through a system of coupled differential equations that simulate both local accumulation and spatial spread.

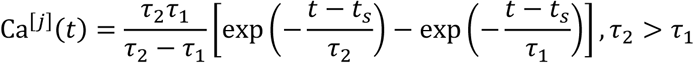

The kinetics are defined by a two-stage process: first, a fast synaptic trace (*x*_Ca_) is triggered by presynaptic input and decays with a time constant *τ*_1_. This trace then feeds into the main calcium pool (Ca), which is regulated by a slower decay constant, *τ*_2_, representing the endogenous buffering and clearance mechanisms of the cell. To account for the spatial heterogeneity of the dendritic tree, the model includes a custom diffusion mechanism. Unlike standard point-neuron models, this allows Ca^2+^ to flow between adjacent compartments based on the concentration gradient. This diffusion (Ca_diff_) is physically constrained: the diffusion coefficient (*α*^[*i*,*j*]^) is proportional to the ratio of the connected compartments *i*, *j* contact lengths (*l*) and their diameters (*d*), ensuring that the model accurately reflects the morphological properties of the neurons being simulated. *κ* is a scaling factor and Ca_tot_ denotes the total Ca^2+^ in the compartment as the summation of the Ca^2+^ of all synapses *S* on that compartment.

#### Multiscale plasticity rules

The model integrates a sophisticated suite of functional and structural plasticity rules that allow the network to undergo long-term learning and reorganization. Functional plasticity is implemented through a Synaptic Tagging and Capture (STC) mechanism, where individual synapses are marked for potential potentiation or depression based on local Ca^2+^ levels. The model implements a non-linear relationship between local calcium concentration (Ca) and the resulting synaptic tag value, which determines the direction of long-term synaptic changes. At low calcium levels (below 0.2 mM), the tag value remains at zero, representing a lack of synaptic modification. Moderate calcium concentrations trigger a negative tag (approaching - 1.0), marking the synapse for Long-Term Depression (LTD). High calcium concentrations trigger a rapid transition to a positive tag (stabilizing at 1.0), marking the synapse for Long-Term Potentiation (LTP). This double threshold-based mechanism (i.e., BCM-like) captures the biphasic nature of calcium-dependent plasticity, ensuring that only activity exceeding specific biological thresholds leads to stable weight adjustments.

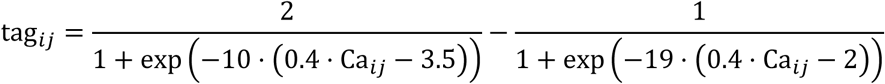

where Ca_*ij*_ denotes the synaptic Ca^2+^ measured in mM.

Once a synaptic tag is set by the calcium-dependent thresholds, its value is not permanent but transient. The temporal evolution of the tag in a synapse from presynaptic neuron *j* to postsynaptic neuron *i* (tag_*ij*_) is governed by a first-order differential equation, causing the tag value to decay exponentially over time.

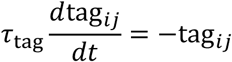

where tag_*ij*_ represents the current state of the synaptic tag (either positive for LTP or negative for LTD). *τ*_tag_ is the tag decay time constant, which determines the duration the synapse remains eligible for permanent strengthening or weakening via PRP capture (see below).

The conversion of a transient synaptic tag into a stable, long-term change is mediated by the synthesis and capture of Plasticity-Related Proteins (PRPs). Our model implements a dual-PRP system triggered by compartmental Ca^2+^ levels. (1) Local PRPs: These are synthesized rapidly following a stimulus, providing an immediate pool of proteins for synaptic consolidation within the local dendritic branch. (2) Global PRPs: These represent a protein synthesis process with a distinct temporal delay. As shown in the model, Global PRPs peak significantly later than Local PRPs, allowing for the reinforcement of synaptic tags over longer intervals. Both PRPs are simulated as alpha functions.

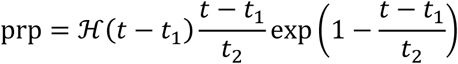

Where for soma *t*_1_ = 20min and *t*_2_ = 30min and for dendrites *t*_1_ = 0min and *t*_2_ = 15min. ℋ(⋅) denotes the Heaviside function:

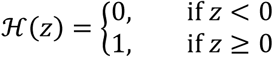

The total availability of these proteins determines whether a decaying synaptic tag is successfully captured and converted into a permanent weight change.

The final change in synaptic strength (Δ*w*_*ij*_) is determined by the convergence of temporal, spatial, and activity-based signals. This “Capture” process is governed by a four-factor rule.

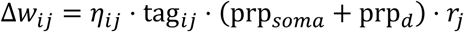

where Δ*w*_*ij*_ represents the total change in the synaptic weight from presynaptic neuron *j* to postsynaptic neuron *i* located onto dendrite *d*, *η*_*ij*_ the synaptic learning rate, which we define as being inversely proportional to the current synaptic strength to ensure stability (see below). tag_*ij*_ is the current value of the synaptic tag. This captures the local calcium history of the synapse, determining if it is eligible for LTP (positive tag) or LTD (negative tag). prp_*soma*_ and prp_*d*_ is the availability of PRPs in the soma and dendrite *d* of the postsynaptic neuron. This acts as a permissive gate; if no proteins are available, the tag cannot be converted into a permanent weight change. *r*_*j*_ denotes the firing rate (activity) of the presynaptic neuron. This ensures that the synapse only updates when there is active communication, preventing weight drift during periods of inactivity (Hebbian-like plasticity). The synaptic learning rate, the scale factor, is a function of the current synaptic weight. It follows a bio-inspired, inversely proportional relationship where weak synapses have a high learning rate, allowing for rapid initial acquisition. In contrast, as synapses strengthen, the scale factor approaches zero, providing homeostatic stability and preventing runaway excitation by slowing the rate of change for established connections.

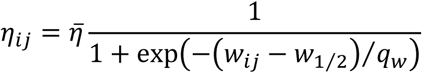

where *η̅* is the maximum asymptotic learning rate, defining the upper bound of the sigmoidal function, *w*_*ij*_ denotes the instantaneous synaptic weight of that specific connection, *w*_1⁄2_ is the half-activation threshold, and *q*_*w*_ is the slope factor that dictates the steepness of the sigmoidal curve.

The model incorporates a dynamic structural plasticity framework known as synaptic turnover, which physically reshapes the network architecture based on ongoing neural activity. This process is driven by two counterbalancing mechanisms: synaptic gain and synaptic loss. Synaptic gain, or formation, is a proactive growth mechanism triggered by high postsynaptic activity. When a specific dendritic compartment exhibits sustained high calcium (Ca^2+^) levels, the model initiates the generation of new synapses. This allows the network to invest structural resources in dendritic branches that are already successfully integrating information, thereby increasing the probability of capturing future relevant inputs in those specific regions. In contrast, the network maintains efficiency and metabolic economy through synaptic loss, or elimination. This pruning process targets connections that are no longer contributing meaningfully to the network’s function. Specifically, the model identifies and removes relatively small synapses with low weights, which are considered “weak” connections that have failed to achieve long-term potentiation through the functional tagging and capture process. Furthermore, synapses that do not receive consistent presynaptic input are also eliminated. By systematically pruning these inactive or redundant pathways, the model reduces noise and prevents overfitting, ensuring that the final circuit architecture remains sparse, reliable, and highly informative.

To further stabilize the network during periods of intense learning and structural reorganization, the model incorporates a synaptic scaling homeostatic mechanism. This rule serves as a global regulator of excitability, ensuring that the total sum of all plastic synaptic weights within the network remains constant over time. While the functional and structural plasticity rules previously described allow individual synapses to grow or shrink in response to specific activity patterns, synaptic scaling works in the background to normalize these changes across the entire population *N*.

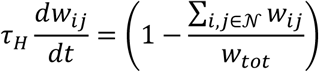

where *τ*_*H*_ denotes the time constant of the scaling process.

By maintaining a fixed total synaptic weight, the mechanism prevents the network from falling into states of pathological activity, such as runaway excitation or total silence. If a group of synapses is significantly strengthened (LTP) due to an associative learning event, the homeostatic rule proportionally scales down the remaining synapses to compensate. This preserves the relative differences in synaptic strength (i.e., which encode the learned information) while ensuring that the overall input drive to the neurons stays within a biologically functional range. This balance is critical for the long-term stability of the recurrent neural network, as it allows for continuous learning without compromising the integrity of the circuit’s baseline dynamics.

#### Rule-specific, contextual, and reward pathways

External inputs to the network are modeled as independent populations of Poisson-spiking afferent neurons, functionally segregated into rule-specific feedforward (FF1, FF2), top-down feedback (FB1, FB2) representing the two modalities, and reward (RWD) pathways. The rule-specific feedforward inputs and the contextual feedback signals each consist of 25 neurons. During the 500 ms stimulus presentation window of each active session, these FF and FB populations fire at a baseline rate of 20 Hz. These afferents structurally target distinct dendritic domains: FF pathways innervate both basal and distal apical compartments, whereas FB pathways project exclusively to the distal tufts of the pyramidal neurons. To mediate reinforcement, the global reward signal is operationalized as a smaller, distinct population of 10 neurons. Because the simulation represents continuous multi-trial sessions rather than discrete choices, the magnitude of the RWD population’s firing rate dynamically scales to represent the overall behavioral accuracy of that specific session. During the final 250 ms of the stimulus period, this performance-scaled RWD rate (reaching a maximum of 50 Hz for the highest accuracy) provides targeted excitatory drive. Mechanistically, these RWD afferents selectively innervate the proximal apical compartments of the pyramidal (PYR) cells and the VIP interneurons, delivering the required depolarization to modulate calcium influx and trigger subsequent weight-dependent structural plasticity.

#### Simulation protocol and network dynamics

The functional performance of the network model was evaluated using a structured simulation protocol spanning 12 distinct sessions and involving two different behavioral rules. Each session consists of a stimulus period with duration 1,000ms (dt=0.1ms) followed by an interstimulus period, where the plasticity mechanisms take place, with duration 24h (dt=1min). This setup replicates experimental paradigms used in studies to observe cognitive flexibility and rule-dependent neural encoding. During the initial phase (Sessions S1-S5), the network was trained under “Rule 1”, characterized by specific patterns of sensory input and reward feedback. As shown in the activity plot, neural firing is precisely organized according to these task demands. The “Rule Switch” occurs between Sessions S5 and S6. Following this transition, the network must adapt its internal representation to accommodate “Rule 2”. To validate the model’s ability to handle cognitive flexibility and long-term stability, we conducted a series of simulations across two experimental groups: a Switch group and a Stable group. Each group consisted of 20 random seeds, which serve as virtual animals representing the variability inherent in biological subjects.

#### Population Firing Rate Analysis

To evaluate the macroscopic activity dynamics of the network during initial learning and subsequent rule-switching, we tracked the mean firing rate of the pyramidal cell population. For each independent network instance (virtual animal or run) and training session, the firing rate (Hz) of individual pyramidal neurons was calculated based on their total spike counts during the session. The overall population firing rate was determined by averaging the activity of all pyramidal neurons within a given run. Temporal trajectories of population activity were subsequently aggregated across all independent runs, with variability expressed as the standard error of the mean (SEM).

#### Identification of active neurons

We operationally defined active neurons using a rigorous, network-specific functional thresholding procedure based on baseline excitability. To account for intrinsic variations between independent network initializations, a baseline activity distribution was established for each run using the firing rates recorded during the initial, naive learning phase (Sessions 1 and 2). For each network, we calculated the baseline mean firing rate (*μ_baseline_*) and its standard deviation (*σ_baseline_*). A global, run-specific activation threshold (*f_θ_*) was then defined to identify significantly elevated activity while preventing the spurious classification of minimally active cells (hard absolute minimum threshold of 1.0 Hz) in networks with exceptionally low baseline excitability. This threshold was mathematically formalized as:

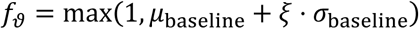

where *ξ* = 1.

Consequently, a pyramidal neuron was classified as active in any given session if, and only if, its firing rate strictly exceeded both its respective global statistical threshold and the absolute 1.0 Hz minimum. The fractional size of the active population was quantified per session by calculating the percentage of pyramidal neurons satisfying these dual criteria.

#### Quantification of structural plasticity (spine dynamics)

We quantified the rates of synaptic gain, loss, and overall turnover using mathematical formulations identical to those employed in the corresponding *in vivo* longitudinal imaging experiments. For each independent network (run) and tracking interval (session *t*), we classified the status of every synapse into one of three mutually exclusive categories: newly formed (*N*_*formed*_), eliminated (*N*_*elim*_), or persistent (*N*_*pers*_). Persistent synapses were defined as those that structurally survived from the previous session (*t* − 1) to the current session (*t*), regardless of changes in their synaptic weights (i.e., encompassing stable, LTP, and LTD synapses). Using these core counts, we defined the total synaptic pool present in the previous session (*S*_*t*−1_) and the total synaptic pool present in the current session (*S*_*t*_) as follows:

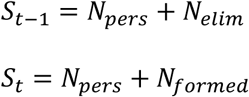

To allow for standardized comparisons across networks with varying absolute numbers of synapses, structural dynamics were calculated as relative percentages.

Spine Gain was defined as the fraction of the current synaptic pool that was newly formed:

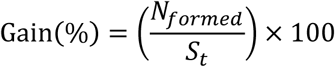

Spine Loss was defined as the fraction of the previous synaptic pool that was eliminated:

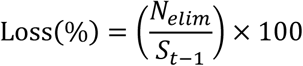

Spine Turnover, representing the total magnitude of structural rewiring, was calculated as the sum of all dynamic events (formation and elimination) divided by the combined total pool of synapses across both timepoints:

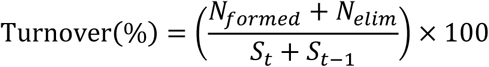

For statistical comparisons of post-switch dynamics (e.g., Stable vs. Switch conditions), these session-by-session metrics were averaged across the entire post-switch learning phase for each independent run to yield a single representative value per metric per network.

#### Calculation of reward-associated synaptic elimination

To quantify the precise timing (exploration vs. learning) of the synaptic elimination, we analyzed the concurrent network reward rate at the precise session of each synaptic elimination event. First, we extracted the mean firing rate of the reward-encoding input population (ℛ) for every session (*t*). To account for baseline differences in reward magnitude across independent network initializations, we normalized the reward rate on a per-run basis. The normalized reward rate, ℛ_*norm*_(*t*), was calculated by dividing the mean reward firing rate of a given session by the maximum reward firing rate observed across all sessions for that specific network run.

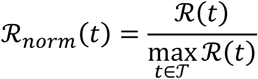

where *T* represents the total set of recorded sessions for the given network. This bounded the reward activity metric between 0 and 1, facilitating direct cross-network comparisons.

Next, eliminated synapses within the Switch condition were strictly stratified into two temporal cohorts based on their developmental origin: ‘Pre-existing synapses’ (spines that existed/formed prior to the rule switch and persisted into the second phase before being eliminated) and ‘Newly formed synapses’ (spines that both formed and were subsequently eliminated entirely within the post-switch learning phase). For every eliminated synapse in both cohorts, we identified its session of elimination (*t*_*loss*_) and assigned it the concurrent normalized reward rate of the network, ℛ_*norm*_(*t*_*loss*_). The resulting distributions of reward rates at the time of synaptic death were aggregated and statistically compared between the newly formed and pre-existing cohorts to determine if one population was preferentially pruned under specific reward contexts.

#### Structural and functional classification of dendrites and neurons

*Dendritic Tagging and Classification*. To map the subcellular distribution of task-specific inputs, individual dendritic compartments were classified based on their local synaptic allocation and long-term connection survival. For each independent network phase (i.e., Rule 1 or Rule 2 exploration and learning), the structural acquisition of novel features was quantified by the net gain of persistent synapses belonging to specific input pathways (e.g., Rule 1 or Rule 2) within each dendrite. This net gain was calculated as the total number of phase-specific synaptic formations minus the total number of phase-specific eliminations for that pathway (bounded at a minimum of zero). To identify dendrites with significant synaptic net gain, we established a dynamic, run-specific allocation threshold defined as the mean net gain plus one standard deviation (*μ*_*net*_ + *ξ* ⋅ *σ*_*net*_, *ξ* = 1). A dendrite was considered active for a given rule if its net gain strictly exceeded this threshold. To determine whether a previously established dendritic representation persisted following the behavioral switch, we implemented an exact synaptic mapping protocol using unique pre- and post-synaptic connection identifiers. A locked historical registry of “legacy” synapses was established at the exact snapshot of the rule switch (*S* = *S*_*switc*ℎ_), capturing all active initial connections alongside additions during Rule 1 presentation. True legacy eviction (*N*_*elim*_) during Rule 2 presentation was isolated by executing a strict connectomic set intersection between post-switch elimination events and this locked registry. This formulation intrinsically discarded ongoing, input-driven pathway turnover noise (synapses both born and pruned entirely within Rule 2 presentation phase). The allowable structural decay threshold (*ϑ*_*elim*_) for each seed was computed natively from the un-inflated legacy eviction distributions (*μ*_*elim*_ + *ξ* ⋅ *σ*_*elim*_, *ξ* = 1). A Rule 1 dendritic representation was tagged “kept” if its verified legacy losses remained strictly below this threshold, indicating the memory trace was successfully protected from catastrophic overwriting. Using these criteria, dendrites were tracked longitudinally and assigned to one of five distinct functional classes at the end of the post-switch phase (End of Rule 2). (1) Rule 1: The dendrite satisfied the criterion for the net gain for Rule 1 synapses during the initial learning phase and successfully protected its legacy pool (met the survival criterion) through the rule switch, without significant Rule 2 feature acquisition; (2) Rule 2: The dendrite failed to acquire Rule 1 during the initial phase, but exhibited significant net formation of Rule 2 synapses during the post-switch phase; (3) Both: The dendrite satisfied both the initial Rule 1 net gain and post-switch survival criteria while concurrently meeting the Rule 2 net gain threshold, indicating localized multi-rule integration; (4) Rule 1 → Rule 2: The dendrite successfully acquired the Rule 1 representation during the Rule 1 learning phase, but subsequently suffered catastrophic forgetting (verified legacy evictions met or exceeded the decay threshold) while concurrently meeting the net gain criteria for Rule 2; and (5) None: The dendrite failed to meet the criteria for either rule during the Rule 2 learning phase (not remaining Rule 1 or Rule 2).

#### Neuronal classification and engram recruitment

The classification of neurons required the convergence of both structural (dendritic) and functional (somatic firing) criteria. First, we evaluated the structural status of each pyramidal neuron by aggregating the tags of its constituent dendrites. A neuron was considered structurally responsive to a given rule if it possessed a minimum percent of dendrites (*θ>=25%* of the available dendrites) classified for that rule (inclusive of dendrites tagged as ‘Both’). Second, to ensure that this structural connectivity translated into functional recruitment, we applied a somatic activity filter. A neuron was considered functionally active if its firing rate during the target session (the final session of Rule 1 or Rule 2) exceeded a baseline-derived threshold (see *Identification of active neurons)*. Neurons satisfying both the structural requirement and the functional somatic requirement for a specific rule were tagged accordingly (Rule 1 or Rule 2). Neurons meeting the dual criteria for both rules were classified as Both. Neuron satisfied the structural and functional criteria for Rule 1 during the initial learning phase, but subsequently failed to maintain Rule 1 status after the switch, while successfully meeting the structural and functional criteria for Rule 2 as Rule 1→ Rule 2. Any neuron that failed to meet the functional firing rate threshold, regardless of its dendritic structural composition, was downgraded and classified as None.

#### Calculation of theoretical chance overlap

To establish a statistical baseline for the integration of task-specific inputs, we calculated the expected theoretical overlap (the ‘Both’ category) assuming that recruitment to Rule 1 and Rule 2 occurred independently and strictly by chance. Because we are evaluating a finite, constant pool of network components (dendrites or neurons) at a specific temporal snapshot (the end of the post-switch phase), the expected random overlap is mathematically modeled by the expected value of a hypergeometric distribution. Let *N*_*total*_ represent the total number of items (either the complete simulated pool of dendrites or the complete pool of pyramidal neurons) analyzed across the network. At the end of the post-switch learning phase, let *N*_*R*_1__ denote the total number of items structurally or functionally recruited by Rule 1 (inclusive of items tagged as ‘Both’), and let *N*_*R*_2__ denotes the total number of items recruited by Rule 2, inclusive of items classified as ‘Both’ or as having transitioned from Rule 1 to Rule 2 (‘Rule 1 → Rule 2’). If recruitment to Rule 2 is modeled as drawing a random sample of size *N*_*R*_2__ without replacement from the total population *N*_*total*_, where the *N*_*_R_*1_ items represent predefined “successes” within that population, the number of overlapping items *X* follows a hypergeometric distribution.

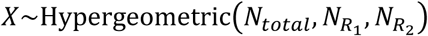

The expected absolute count of randomly overlapping items, *E*[*X*], is mathematically defined as the product of the sample size and the proportion of successes in the total population.

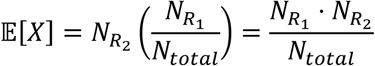

To map this expected theoretical count to our normalized network metrics, we converted it into an expected percentage. The theoretical chance level (%_*c*ℎ*ance*_) plotted alongside the empirical network data was calculated as the expected overlap count divided by the total population pool.

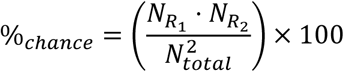

This baseline formulation ensures that the observed fractions of the ‘Both’ functional classes can be rigorously compared against the exact mathematical expectation of independent, random recruitment at the specific timepoint of evaluation.

#### In silico ablation of subcellular integration

To mechanically isolate the computational role of dendritic overlap and spatial clustering during the rule-switching paradigm, we designed two in silico targeted structural constraints. These conditions altered the spatial distribution rules of the network to prevent the integration of task-specific inputs. In the “No Overlap” condition, we specifically disrupted the capacity for local synaptic integration between the sequential tasks. Following the initial learning phase, any dendritic compartment that successfully met the classification criteria for a Rule 1 representation was artificially restricted from receiving newly formed Rule 2 synapses. Consequently, Rule 2 inputs were forced to innervate naïve dendritic branches, completely eliminating the possibility of subsequent subcellular overlap (thereby preventing the formation of ‘Both’ dendrites). To test the extreme functional boundary of spatial separation, we implemented a strict, predefined segregation protocol. In this network variant, the synaptic formation rules were explicitly constrained such that all synapses associated with Rule 1 (including both feedforward and feedback pathways) and all synapses associated with Rule 2 were forced to instantiate on completely distinct, non-overlapping subsets of dendritic compartments. This condition guarantees the physical segregation of the two task representations across the dendritic arbor from the outset of the simulation. However, this criterion also reduces -by half- the pool of available dendrites that each rule can occupy, forcing the potential memory engram into a much smaller set of dendrites. To ensure unbiased functional comparisons despite this artificial reduction in the available dendritic pool, the structural recruitment threshold for each neuron (i.e., the minimum number of rule-specific branches required for engram classification) was scaled to 25% of the neuron’s available dendritic pool.

### Statistics and reproducibility

All procedures were approved by the Veterinary Directorate of the Prefecture of Heraklion (Crete) and the FORTH Ethics Committee. The imaging cohort comprised four Switch-group mice (28 dendrites, 645 spines) and four Stable-group mice (15 dendrites, 358 spines); the pharmacology cohort comprised six saline- and six muscimol-treated mice. Mice were randomly allocated to groups and, in the pharmacology experiment, randomized to receive saline or muscimol. No animals were excluded from the behavioral analyses. Dendritic segments were included only when image quality was maintained across all sessions (low background, isolation from neighboring fluorescent structures, and healthy morphology without marked density loss, smoothing or blebbing); mice whose imaging quality degraded substantially were excluded, and no statistical outliers were removed.

Data distributions were assessed for normality with the Shapiro–Wilk test; normally distributed data were compared with two-tailed Student’s t-tests and non-normally distributed data with Mann–Whitney U tests. Two-way ANOVA, Pearson correlation, linear regression and Kolmogorov–Smirnov tests were used as indicated in the figure legends and in **Supplementary Table 1**. Session-by-session comparisons between Saline and Muscimol cohorts (Sessions 0-9) were evaluated using two-sided Mann-Whitney U test. To quantify phase errors, sessions were split into Early (Sessions 0-4) and Late (Sessions 5-9) phases. Between-group differences within each phase were tested using Welch’s two-sample t-tests, while longitudinal changes within cohorts were evaluated using two-tailed paired t-tests and multiple comparisons across phase-level tests were adjusted using Bonferroni correction (k = 4). For spine-clustering analyses, values were aggregated hierarchically —averaged across dendrites within each mouse and then across mice— so that the mouse was the unit of analysis; permutation-based null distributions (1,000 iterations for turnover domain detection, 10,000 for spine clustering) were generated by reassigning spine positions among biologically valid locations while conserving spine density. Data are presented as mean ± standard deviation (SD) unless otherwise indicated. p<0.05 was considered significant (*p<0.05, **p<0.01, ***p<0.001; n.s., not significant); exact p values, tests and sample sizes are reported in **Supplementary Table 1**. Analyses used custom Python (version 3.12) code, and the network model was implemented in brian2 (version 2.8.0) simulator using the Dendrify framework (version 2.2.0). All simulations were run on a high-performance computing (HPC) cluster using the Rocks distribution^67^. Raw data are available from the corresponding author upon reasonable request. The analysis and modeling code will be made publicly available on GitHub and Zenodo following publication.

### Data availability

The raw datasets generated and/or analyzed during the current study and the code will be publicly available upon publication.

## Supplementary Material

**Supplementary Table 1.**
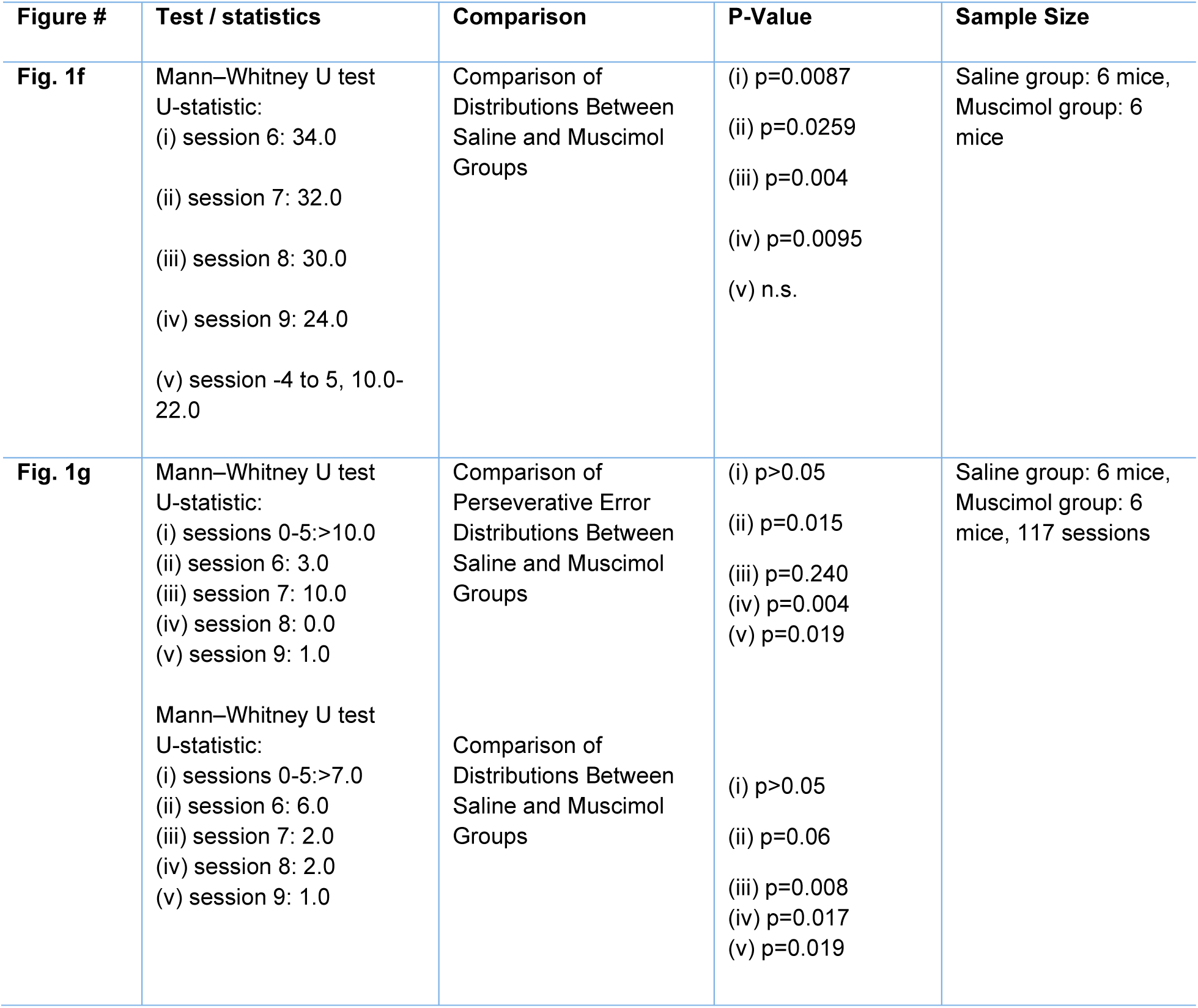
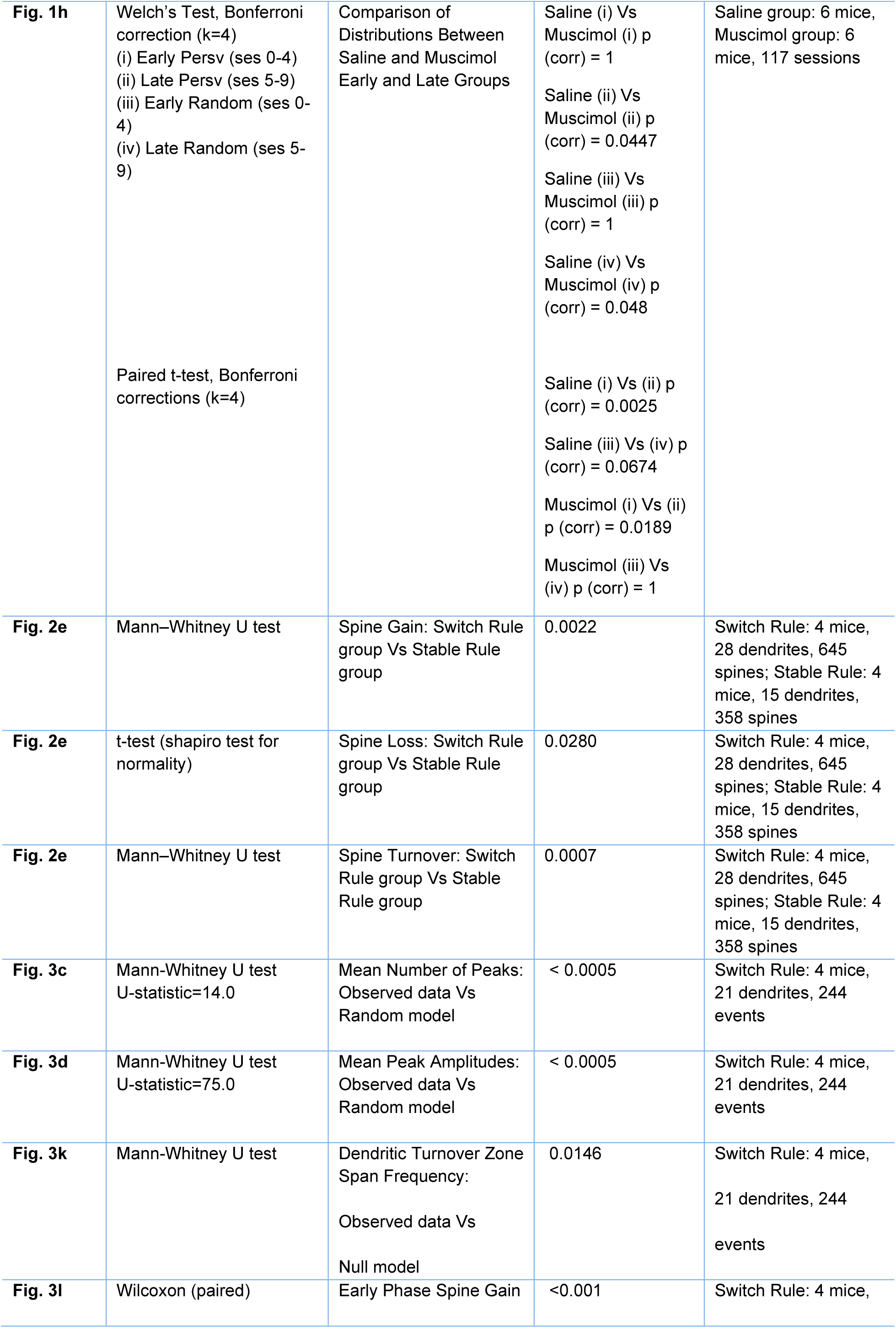
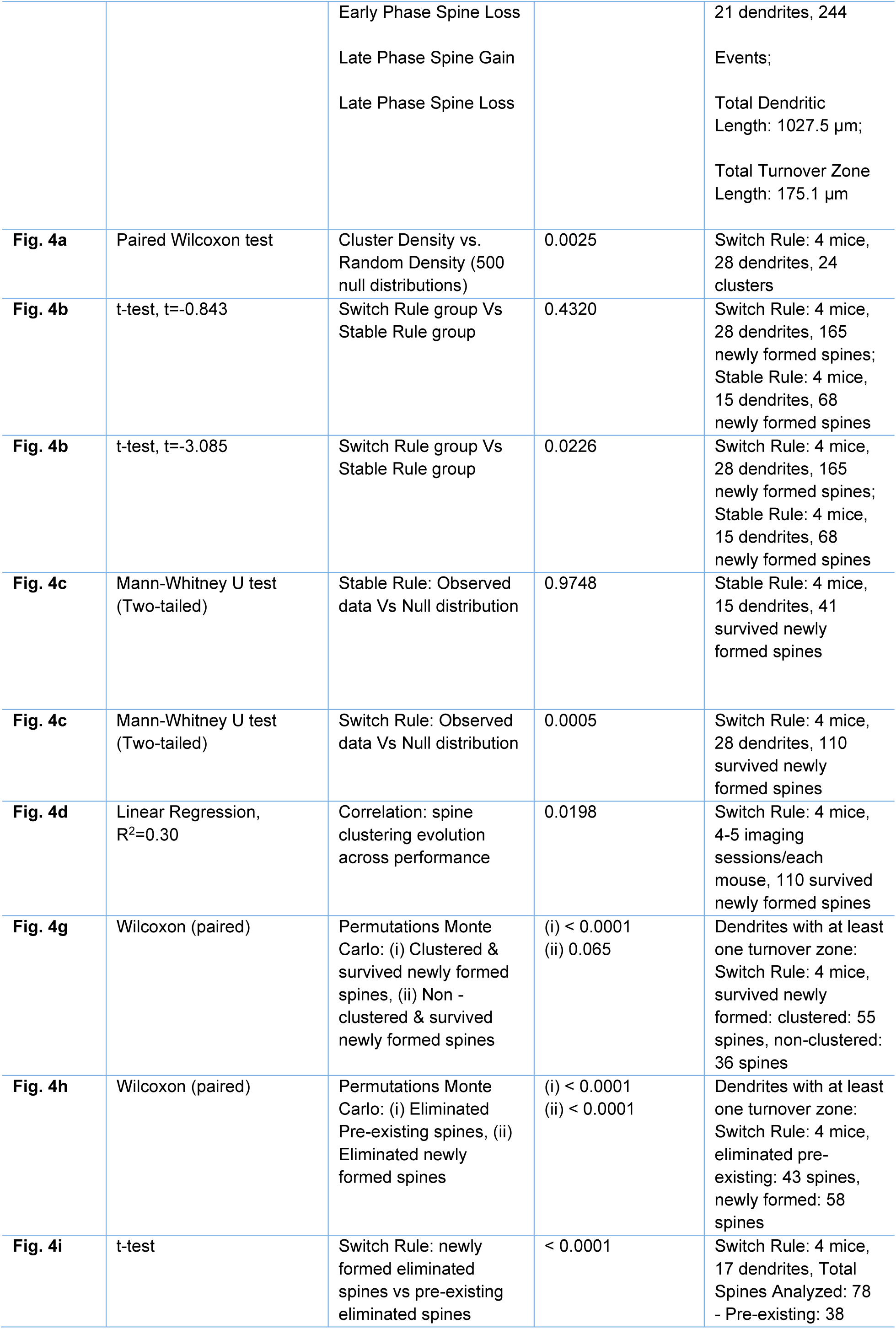
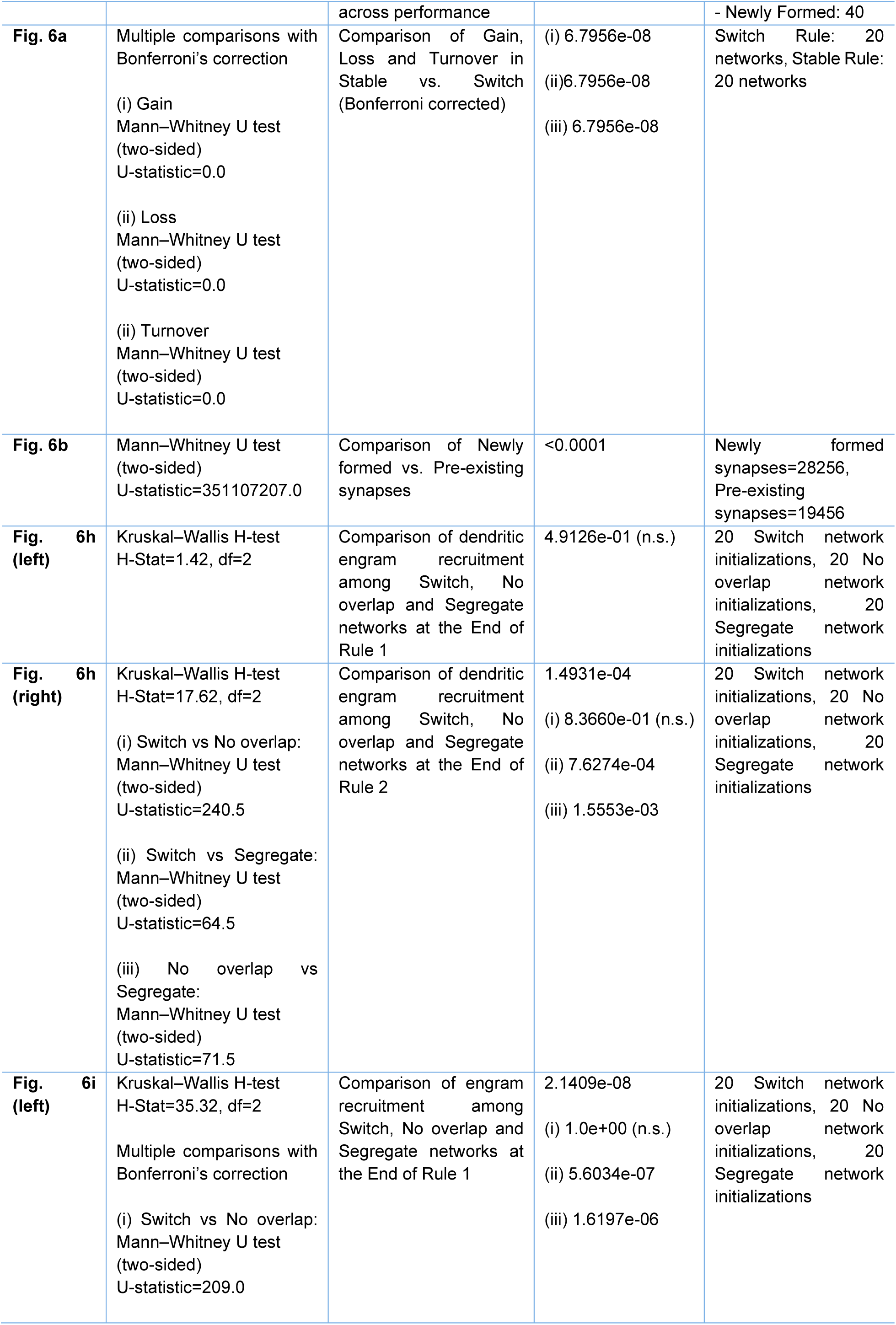
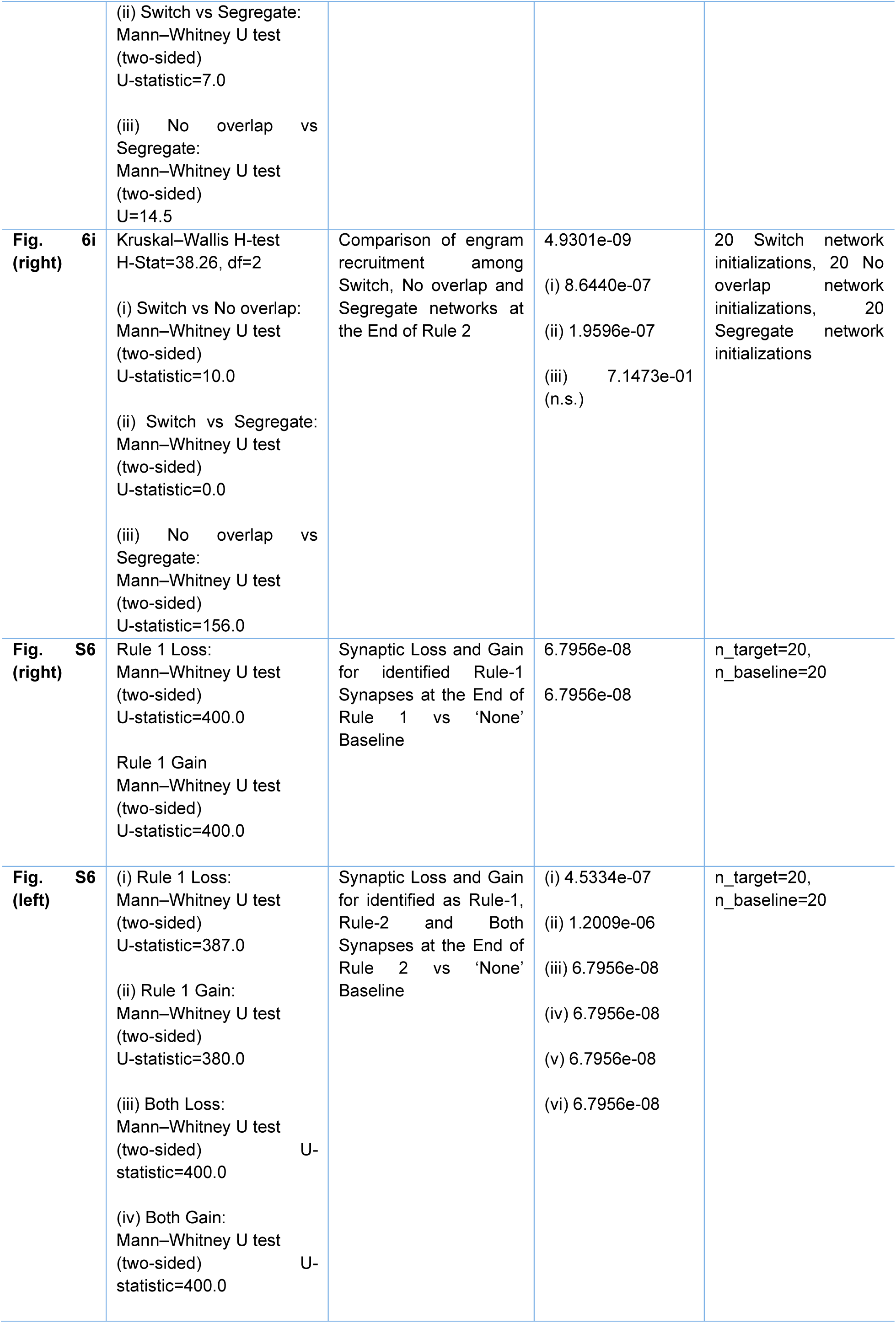
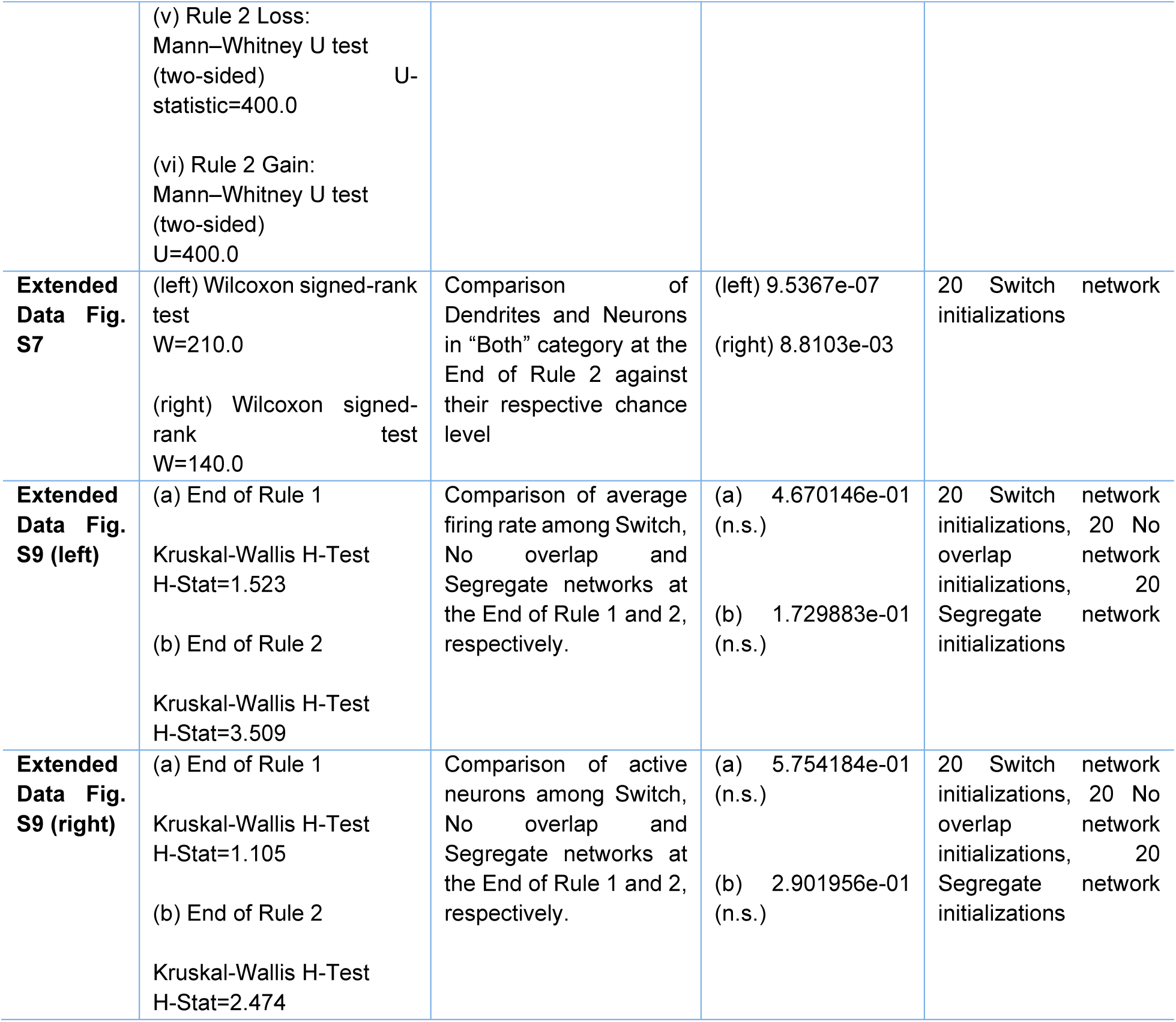
P-values and sample size for all statistical comparisons.

## Extended Figures

**Extended figure S1.**
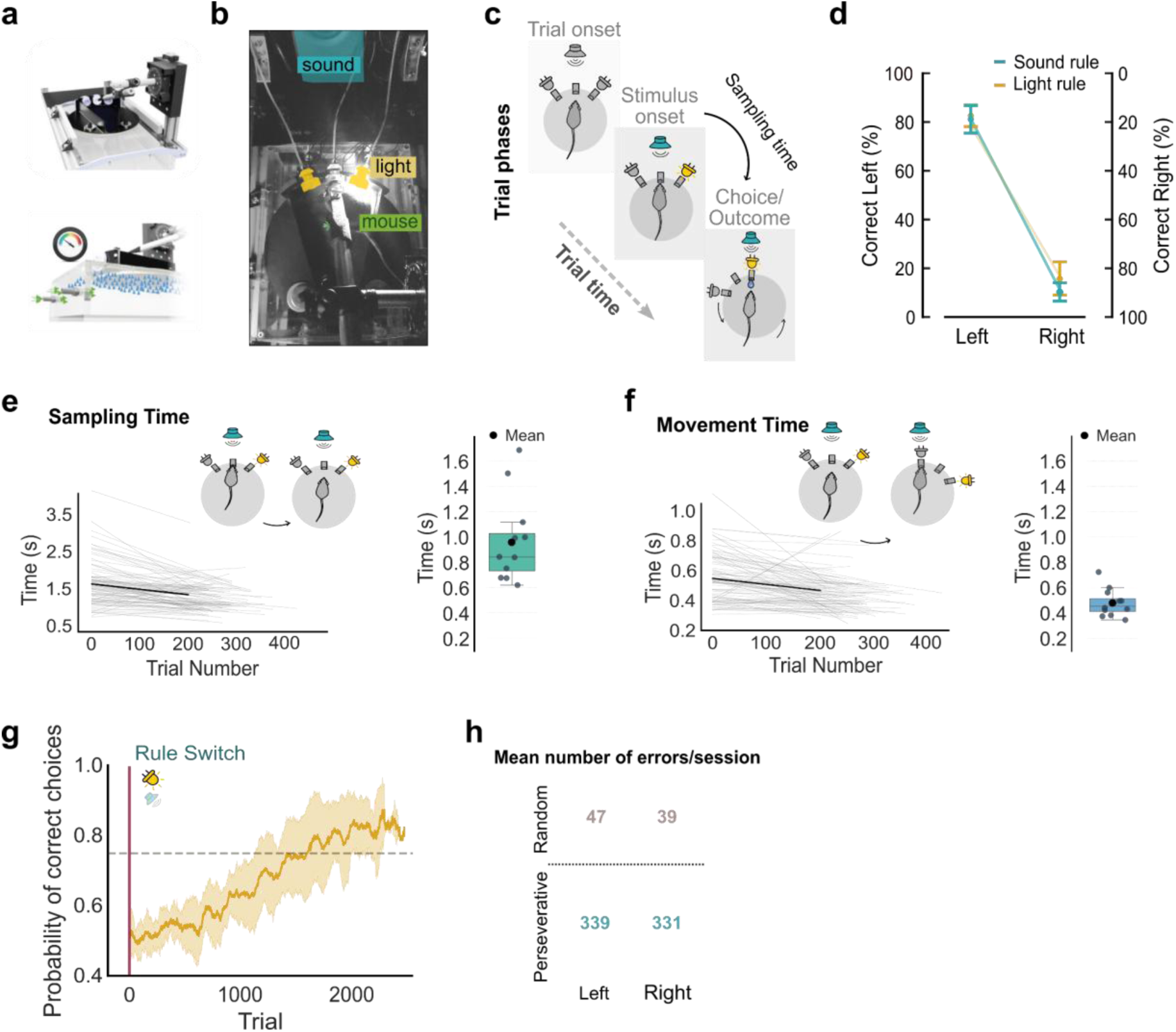
The new rule switch task: a head-fixed, two-alternative forced choice (2AFC) behavioral paradigm. **a**, An illustration of the behavioral apparatus of the task. **b**, The Plexiglas air-table, the floating platform, the head-fixed mice and the rule-specific cues (light and sound) are depicted. **c**, Left: The three sequential phases of a complete trial: Trial onset, Stimulus onset and Choice/Outcome. Stimulus sampling time is measured as the interval from the stimulus onset until the mouse makes a decision by poking into one of the two choice ports; Right: Distribution of the mean reaction time across sessions per mouse. **d**, Psychometric curves: n=11. Mann-Whitney U Test, Correct Left (%): U-statistic=78.0, p=0.2638, Correct Right (%): U-statistic=33.0, p=0.0759. **e**, Stimulus sampling time. Left: Each gray line represents the regression of sampling times for an individual session; the black line indicates the mean regression across all sessions from all mice; n=11 mice. Right: Distribution of the mean sampling time across sessions per mouse. Each dot corresponds to a single mouse (n=11 mice, 98 sessions). Mean sampling time: 0.97s; median sampling time: 0.84s. **f**, Movement Time. Left: Movement time is defined as the duration from the poke out of the center port until the poke in to the choice port. Movement time across trials for multiple sessions. Each gray line represents the regression of movement times for an individual session; the black line indicates the mean regression across all sessions from all mice (n=11 mice). Right: Distribution of the mean movement time across sessions per mouse. Each dot corresponds to a single mouse (n=11, 98 sessions). Mean movement time: 0.48s; median movement time: 0.44s. **g**, Learning Curve. Mean performance across trials calculated using an 80-trial moving window. The dashed line marks the 75% performance threshold. The vertical pink line indicates the rule switch. Mean trial that passes the criterion is 1589; n=11 mice. **h**, Average of total perseverative and random errors after the rule switch for the left and right choice ports. In (**g**) and (**h**) the solid gold line represents the mean performance across all trials and sessions respectively, while the shaded area indicates ±1 SD.

**Extended figure S2.**
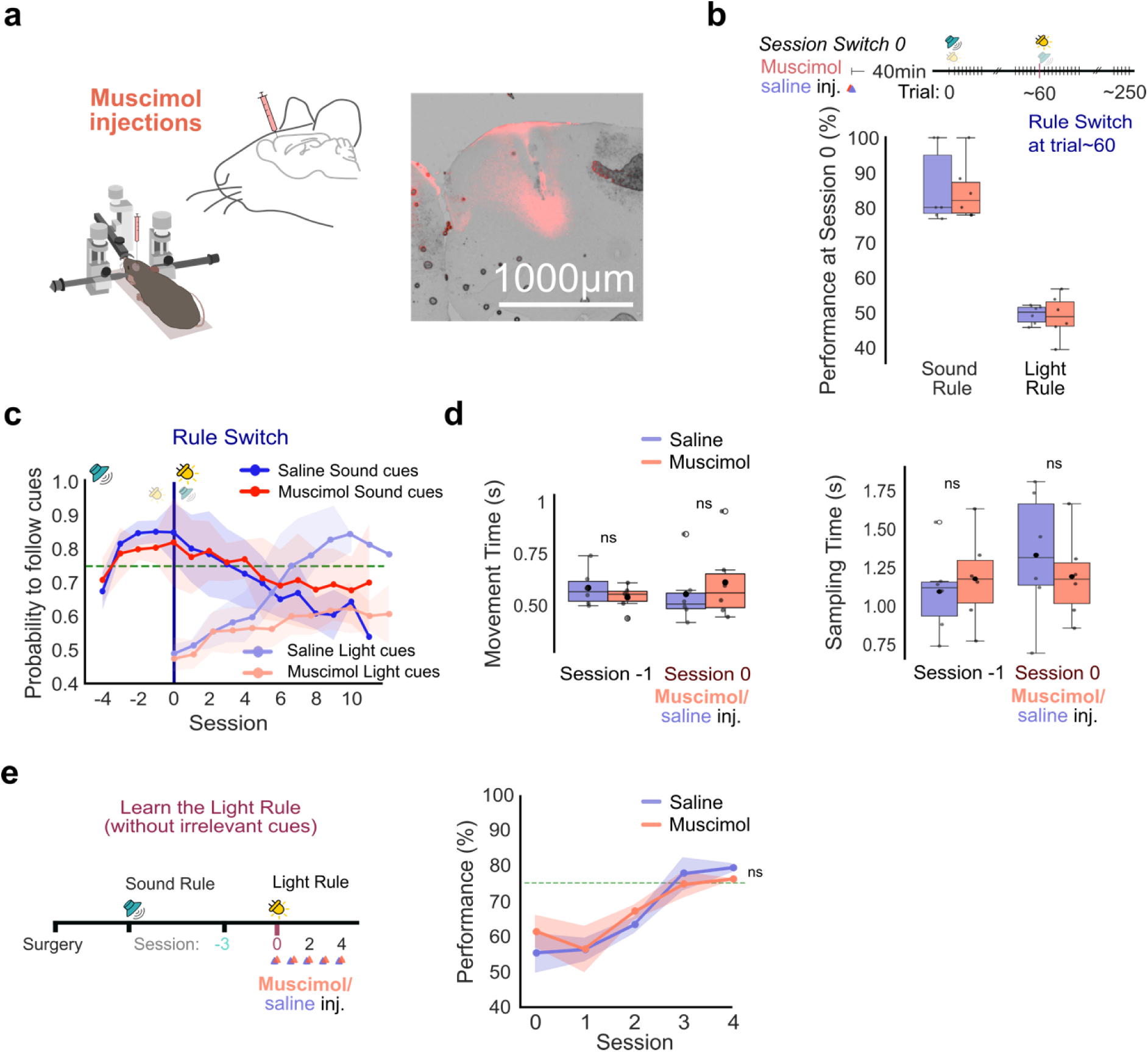
**a**, **Stereotaxic injection of muscimol into the M2 mouse brain area.** Right: Representative fixed brain slice illustrating the diffusion of fluorescent muscimol; histological analysis confirms that the spread is restricted to the cortical layers and does not extend into subcortical structures. Image: bright light and BODIPY TMR-X conjugate. This TMR-X version emits a bright orange/red light (Excitation: ∼ 544 nm / Emission ∼570 nm). Scale bar: 1000μm. **b**, M2 inactivation does not affect performance on the previously learned SR. By switching the rule mid-session (at trial ∼60 of session 0) we separated pre-switch execution (SR) from post-switch updating (LR) within the same animals. blue=saline, red=muscimol groups (n=6 per group; Mann–Whitney U test, SR: U=17.0, p=0.93; LR: U=19.0, p=0.94). **c**, Probability to follow sound/light cues in Saline and Muscimol groups of mice. Following the rule switch, mice initially continue to follow sound cues for several sessions. **d**, M2 silencing does not introduce motor deficits. Analysis of key movement and sampling metrics shows no significant differences between the muscimol and control groups (Saline group: 6 mice, Muscimol group: 6 mice, Movement Time, session -1: p=0.5887, session 0: p=0.5887, Sampling Time, session -1: p=0.4848, session 0: p=0.5887; Mann-Whitney U test). **e**, The performance deficit is specific to sensory conflict. In a control experiment where the conflicting sound cues were removed, M2 inactivation did not prevent mice from successfully learning the LR (Saline group: 3 mice, Muscimol group: 3 mice, mixed ANOVA test: p=0.457). In **b**, **c**, **e** the shaded area indicates ±1 SD.

**Extended figure S3.**
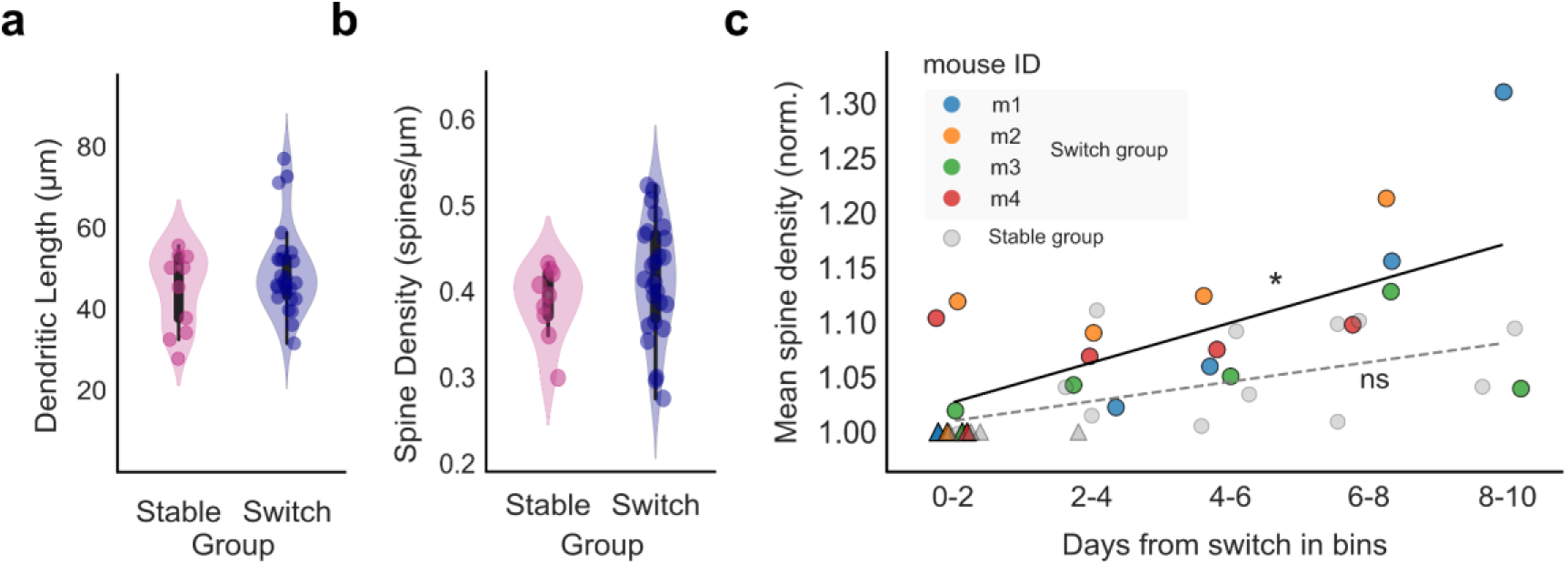
**a**, Average dendritic length and **b**, spine density of dendritic compartments in Stable and Switch Rule group (Switch Rule: 4 mice, 28 dendrites; Stable Rule: 4 mice, 15 dendrites; Dendritic length: statistical test: Mann-Whitney U test: p=0.599, Spine Density: statistical test: t-test: p=0.8413). **c**, Spine density in Switch and Stable Group animals across days following the switch (Switch Rule: 4 mice, Pearson Correlation: r=0.494, p=0.0440; Stable Rule: 4 mice, r=0.234, p=0.0879).

**Extended figure S4.**
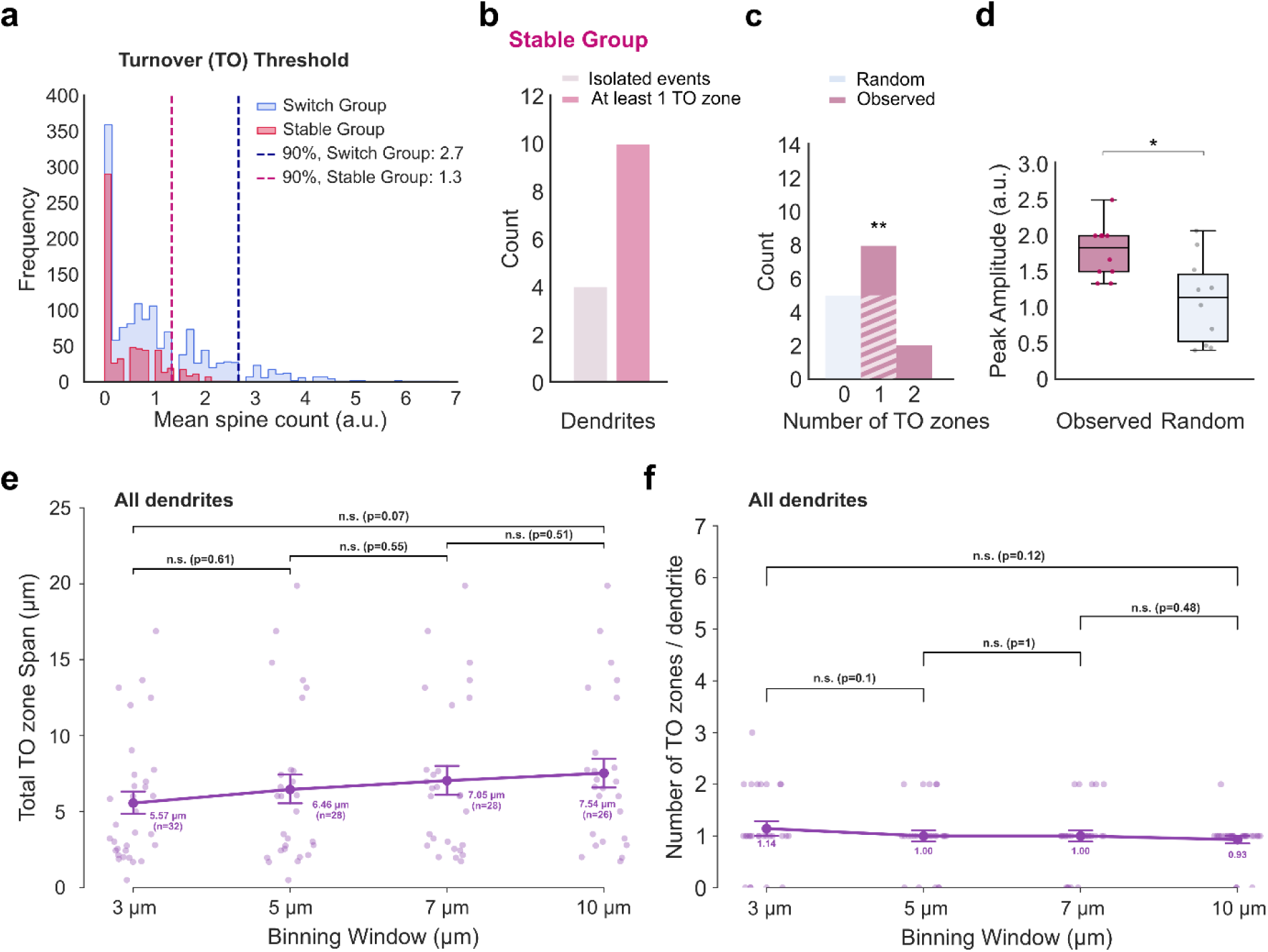
Local density thresholding and validation of spine turnover zones in Stable Group. **a,** Methodological determination of turnover zone thresholds. Representative local event density distributions (spine events per bin) are shown for the Switch Rule (light blue) and Stable Rule (light red) groups. Dashed vertical lines indicate the group-specific 90th percentile thresholds used to define dendritic turnover zones. **b**, Stable Group: The majority of dendritic segments (10 out of 15) have at least one turnover zone. **c**, Stable Group: Comparison of the number of these zones per segment between a randomly shuffled model and observed data: the dendritic compartments where zones were found (Observed: 10 dendrites, 4 mice; Random: 10 dendrites, 4 mice; p=0.0082, Wilcoxon signed-rank test). (**d**), Mean amplitude of binned dynamic events for the random model and the observed data (p=0.014, Paired t-test). (**e**), Parametric exploration of turnover zones physical span across different binning windows (3, 5, 7, and 10 µm) evaluated across all 28 dendritic segments. Turnover zone group span was robust and invariant to window size, with all sequential and endpoint pairwise comparisons being non-significant (p > 0.05, Mann-Whitney U test; large dots indicate mean ± s.e.m., small dots indicate individual turnover zones; n indicate the total number of detected zones). (**f**), Parametric exploration of turnover zone detection frequency across all 28 dendritic segments. The number of detected zones per dendritic segment remained stable across all tested binning window sizes, with all pairwise window comparisons being non-significant (p > 0.05, Mann-Whitney U test; large dots indicate mean ± s.e.m., small dots show individual dendritic segments).

**Extended figure S5.**
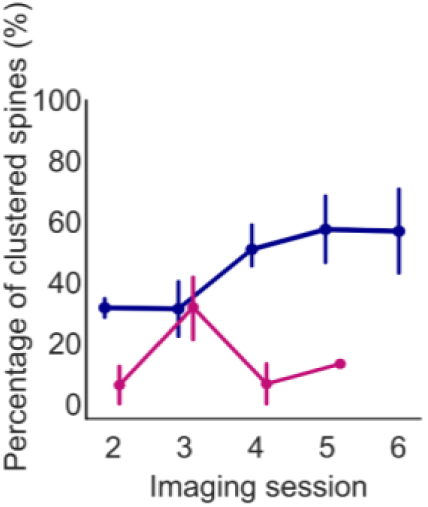
Proportion of clustered new spines across imaging sessions. The proportion of clustered new spines in the Switch group (n=4 mice, deep blue line) scaled across imaging sessions, reaching approximately 60%. In contrast, no such scaling was observed in the Stable group (n=4 mice, magenta line), where performance remained high throughout the imaged sessions. Imaging session 1 corresponds to behavioral session 0 (Rule Switch).

**Extended figure S6.**
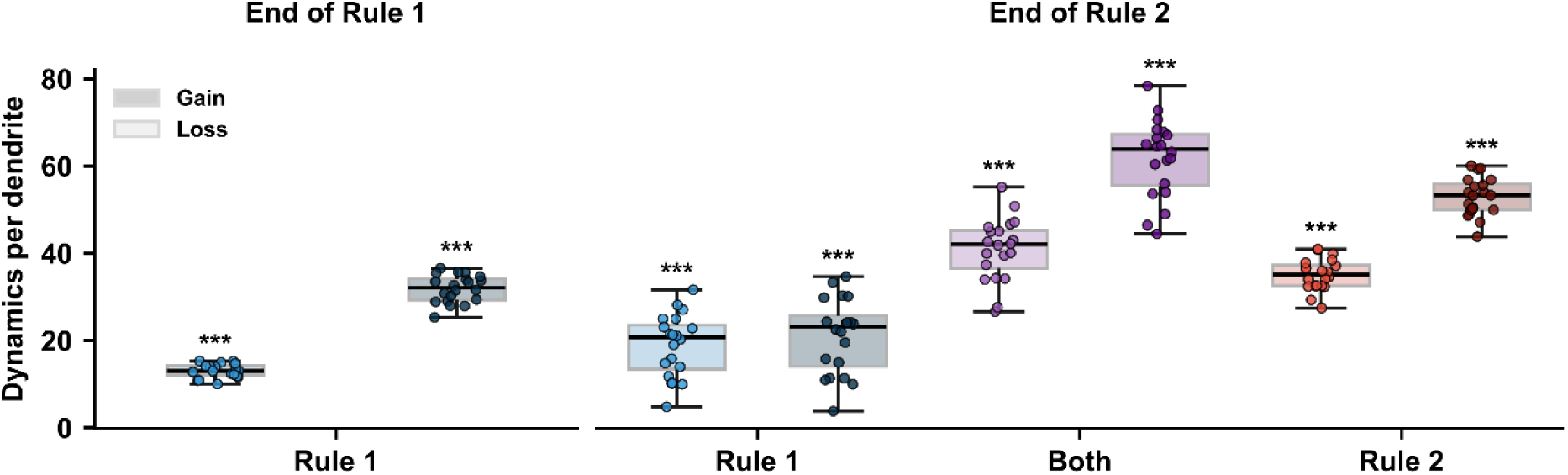
Structural dynamics are elevated in functionally classified dendrites and highest in mixed-selective branches. Spine gain (darker) and loss (lighter) per dendrite by functional class (Rule 1, Both, Rule 2) at End of Rule 1 and End of Rule 2; at the end of Rule 1 only Rule 1-selective dendrites are present. Stars denote dynamics significantly elevated above the unclassified (‘None’) dendritic baseline (Mann–Whitney U test, ***p<0.001). Box plots show median and interquartile range; whiskers extend to minimum and maximum values.

**Extended Figure S7.**
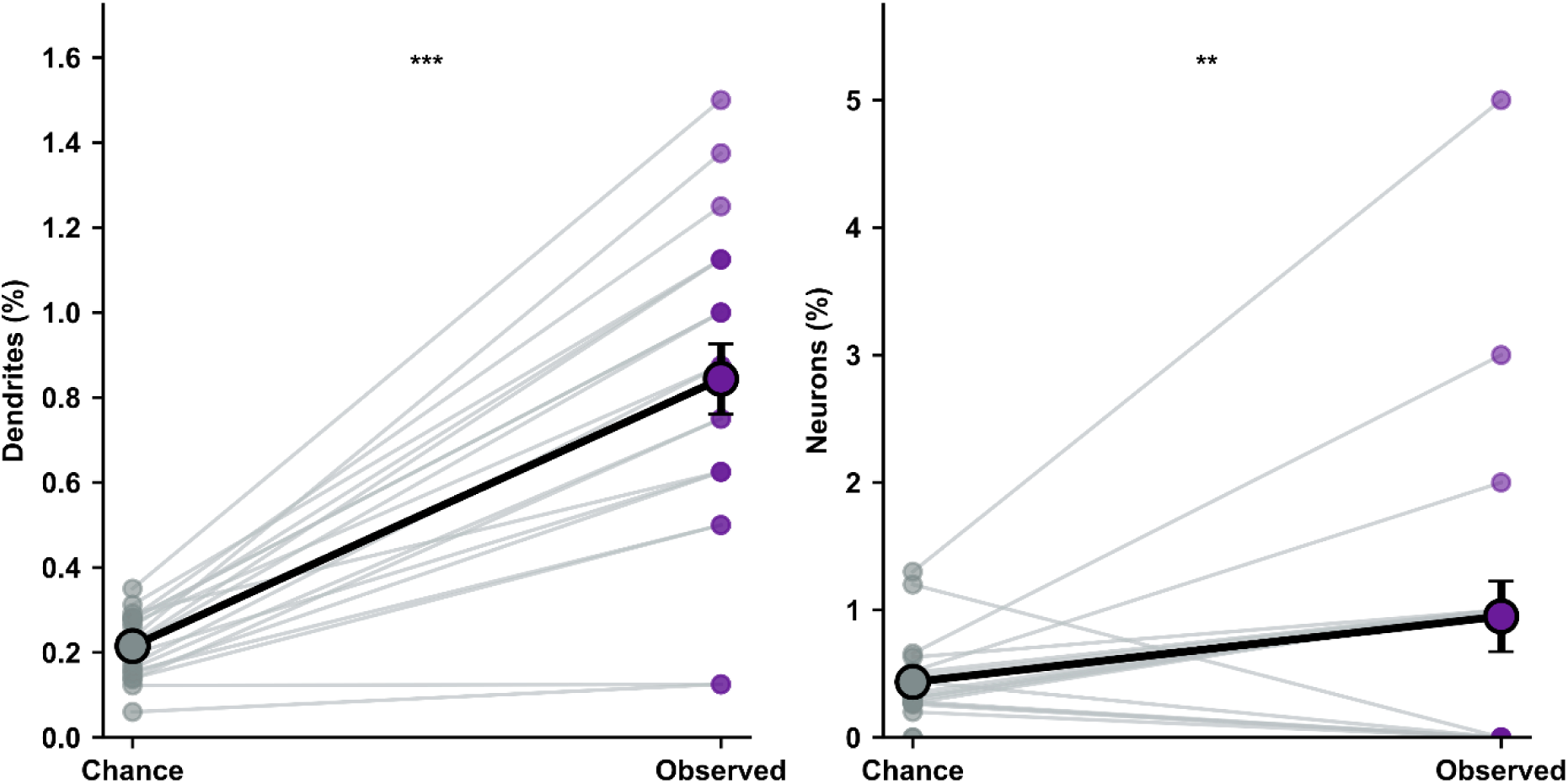
Distribution of multi-rule engram co-allocation against independent analytical baselines. Left, Percentage of dendritic compartments concurrently allocated to both rule representations (Both, purple dots) compared to the run-matched independent analytical chance baseline (Chance, gray dots) at the end of Phase 2. Light gray lines connect the paired Chance and Observed values from each individual simulation run (n=20); black-outlined circles and error bars denote the group mean ± SEM, respectively. Right, Percentage of pyramidal neurons concurrently participating in both active functional engrams, plotted as in the left panel (n=20 independent simulation runs). Stars denote statistical significance (one-sided Wilcoxon signed-rank test; *p<0.05, **p<0.01).

**Extended figure S8.**
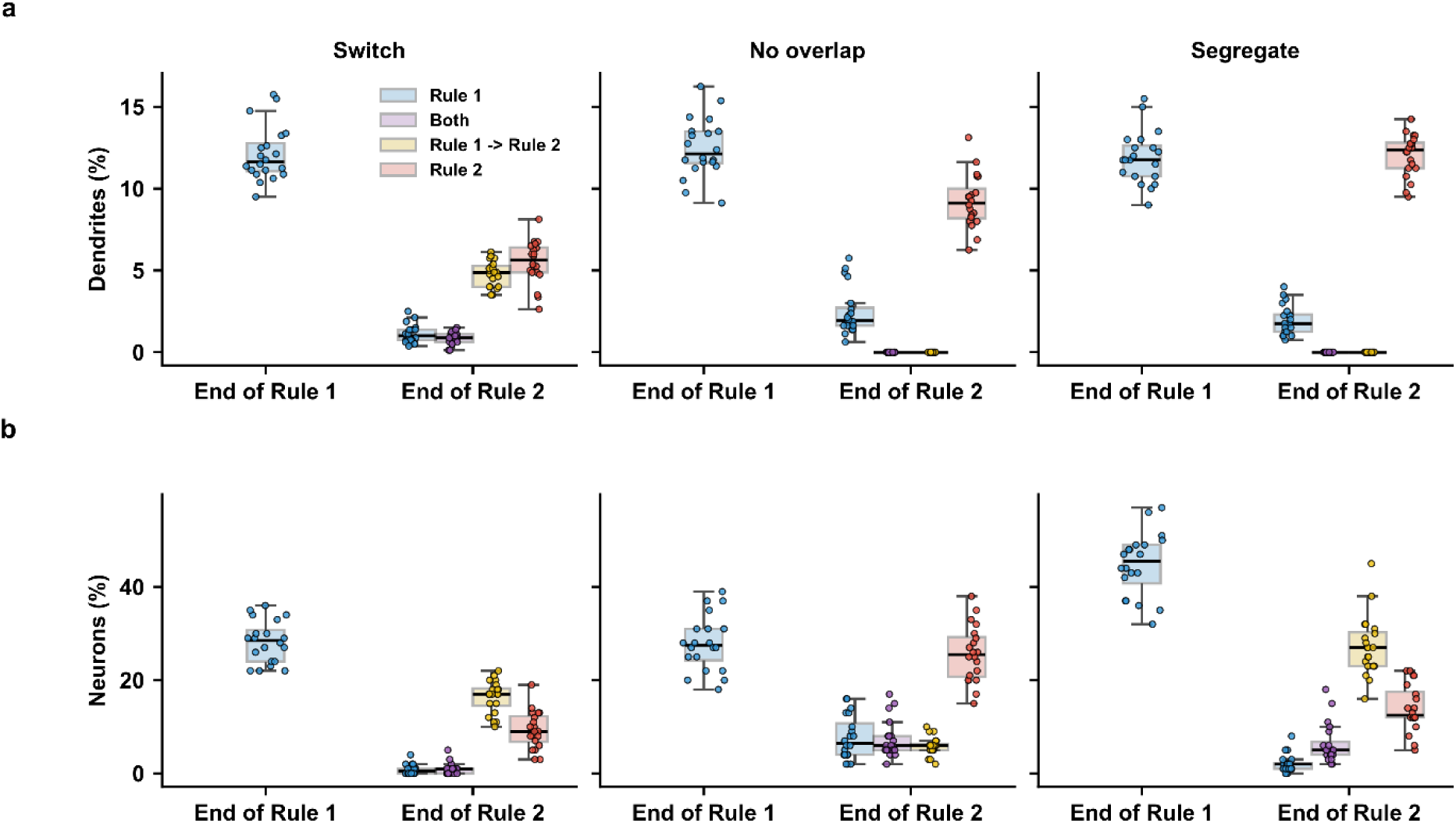
The reuse constraints remove mixed selectivity at the dendritic but not at the somatic level. **a**, Percentage of dendrites classified as Rule 1-selective (blue), Both (purple), transitioning Rule 1 → Rule 2 (yellow) or Rule 2-selective (red) by the combined structural and functional criteria used in Fig. 6f (*Methods*), at the End of Rule 1 (left) and End of Rule 2 (right), shown for the intact Switch network, the No-overlap condition and the Segregate condition. Both constraints abolish mixed-selective and transitioning dendrites by the end of Rule 2, confirming that the manipulations acted as intended. **b**, As in (**a**), for neurons using the same criteria as in Fig. 6g. Mixed-selective and transitioning neurons persist —and mixed-selective neurons become more frequent than in the intact network— under both constraints, indicating that when compartment reuse is blocked the two rules are combined across separate branches of the same neuron rather than being allocated to separate cells. Data for the Switch network in (**a**) and (**b**) are the same as in Fig. 6f, g and are reproduced here for comparison. Box plots show median and interquartile range; whiskers extend to minimum and maximum values.

**Extended figure S9.**
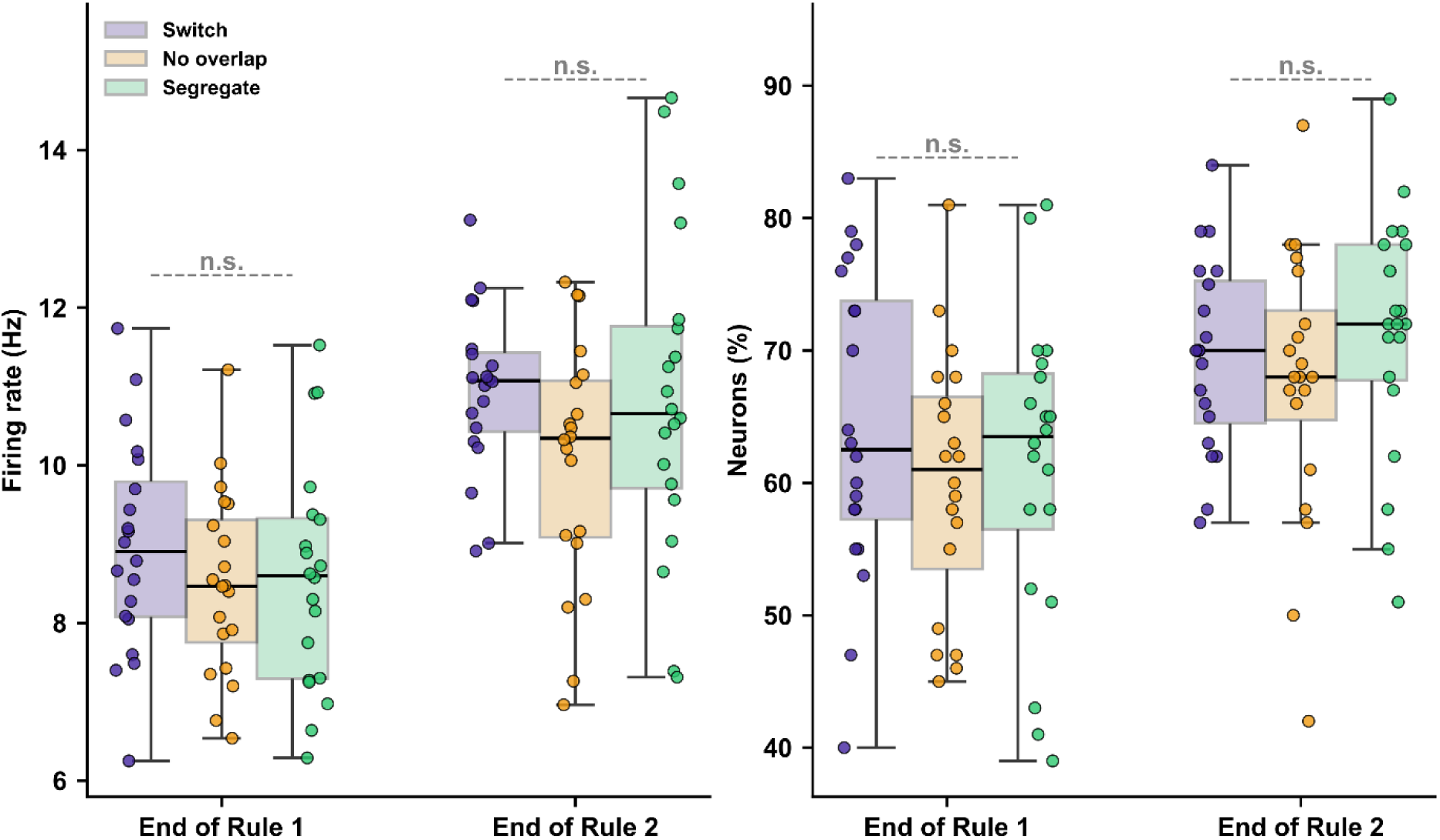
Validation and network stability of the ablated models. Left, mean firing rate (Hz) of all pyramidal neurons for the Switch (purple), No overlap (yellow), and Segregate (green) conditions, evaluated at the end of Rule 1 and the end of Rule 2. Right, Percentage of active neurons, as defined in *Identification of active neurons* section, across the same conditions and timepoints. Boxplots indicate the median and interquartile range, with whiskers extending to the minimum and maximum values. Statistical significance across the three experimental conditions was evaluated at each timepoint using a non-parametric omnibus Kruskal–Wallis H-test; dashed gray lines indicate the omnibus result for that timepoint. Where the omnibus test was significant (p<0.05), pairwise post-hoc Mann-Whitney U tests with Bonferroni correction (m=3 comparisons) were performed between condition pairs, shown as solid black brackets. Significance levels are indicated as labeled (*p<0.05; n.s., not significant, p>0.05).

## References

1. Kirkpatrick, J. et al. Overcoming catastrophic forgetting in neural networks. Proc. Natl. Acad. Sci. 114, 3521–3526 (2017).

2. Holtmaat, A. & Svoboda, K. Experience-dependent structural synaptic plasticity in the mammalian brain. Nat. Rev. Neurosci. 10, 647–58 (2009).

3. Kasai, H., Fukuda, M., Watanabe, S., Hayashi-Takagi, A. & Noguchi, J. Structural dynamics of dendritic spines in memory and cognition. Trends Neurosci. 33, 121–129 (2010).

4. Fu, M., Yu, X., Lu, J. & Zuo, Y. Repetitive motor learning induces coordinated formation of clustered dendritic spines in vivo. Nature 483, 92–95 (2012).

5. Hedrick, N. G. et al. Learning binds new inputs into functional synaptic clusters via spinogenesis. Nat. Neurosci. 25, 726–737 (2022).

6. Lai, C. S. W., Franke, T. F. & Gan, W.-B. Opposite effects of fear conditioning and extinction on dendritic spine remodelling. Nature 483, 87–91 (2012).

7. Lendvai, B., Stern, E. A., Chen, B. & Svoboda, K. Experience-dependent plasticity of dendritic spines in the developing rat barrel cortex in vivo. Nature 404, 876–881 (2000).

8. Xu, T. et al. Rapid formation and selective stabilization of synapses for enduring motor memories. Nature 462, 915–919 (2009).

9. Yang, G., Pan, F. & Gan, W.-B. Stably maintained dendritic spines are associated with lifelong memories. Nature 462, 920–924 (2009).

10. Cichon, J. & Gan, W.-B. Branch-specific dendritic Ca2+ spikes cause persistent synaptic plasticity. Nature 520, 180–185 (2015).

11. Frank, A. C. et al. Hotspots of dendritic spine turnover facilitate clustered spine addition and learning and memory. Nat. Commun. 9, 422 (2018).

12. Palmer, L. M. et al. NMDA spikes enhance action potential generation during sensory input. Nat. Neurosci. 17, 383–390 (2014).

13. Sehgal, M. et al. Compartmentalized dendritic plasticity in the mouse retrosplenial cortex links contextual memories formed close in time. Nat. Neurosci. 28, 602–615 (2025).

14. Chater, T. E., Eggl, M. F., Goda, Y. & Tchumatchenko, T. Competitive processes shape multi-synapse plasticity along dendritic segments. Nat. Commun. 15, 7572 (2024).

15. Kerlin, A. et al. Functional clustering of dendritic activity during decision-making. eLife 8, e46966 (2019).

16. Harvey, C. D. & Svoboda, K. Locally dynamic synaptic learning rules in pyramidal neuron dendrites. Nature 450, 1195–1200 (2007).

17. Harvey, C. D., Yasuda, R., Zhong, H. & Svoboda, K. The spread of Ras activity triggered by activation of a single dendritic spine. Science 321, 136–40 (2008).

18. Wilson, D. E., Whitney, D. E., Scholl, B. & Fitzpatrick, D. Orientation selectivity and the functional clustering of synaptic inputs in primary visual cortex. Nat. Neurosci. 19, 1003– 1009 (2016).

19. Poirazi, P., Brannon, T. & Mel, B. W. Pyramidal Neuron as Two-Layer Neural Network. Neuron 37, 989–999 (2003).

20. Poirazi, P., Brannon, T. & Mel, B. W. Arithmetic of Subthreshold Synaptic Summation in a Model CA1 Pyramidal Cell. Neuron 37, 977–987 (2003).

21. Poirazi, P. & Papoutsi, A. Illuminating dendritic function with computational models. Nat. Rev. Neurosci. 21, 303–321 (2020).

22. Hwang, F.-J. et al. Motor learning selectively strengthens cortical and striatal synapses of motor engram neurons. Neuron 110, 2790–2801.e5 (2022).

23. Wright, W. J., Hedrick, N. G. & Komiyama, T. Distinct synaptic plasticity rules operate across dendritic compartments in vivo during learning. Science 388, 322–328 (2025).

24. Branco, T. & Häusser, M. The single dendritic branch as a fundamental functional unit in the nervous system. Curr. Opin. Neurobiol. 20, 494–502 (2010).

25. Choucry, A., Nomoto, M. & Inokuchi, K. Engram mechanisms of memory linking and identity. Nat. Rev. Neurosci. 25, 375–392 (2024).

26. Kastellakis, G. & Poirazi, P. Synaptic Clustering and Memory Formation. Front. Mol. Neurosci. 12, 300 (2019).

27. Kastellakis, G., Silva, A. J. & Poirazi, P. Linking Memories across Time via Neuronal and Dendritic Overlaps in Model Neurons with Active Dendrites. Cell Rep. 17, 1491–1504 (2016).

28. Hayashi-Takagi, A. et al. Labelling and optical erasure of synaptic memory traces in the motor cortex. Nature 525, 333–338 (2015).

29. Cai, D. J. et al. A shared neural ensemble links distinct contextual memories encoded close in time. Nature 534, 115–118 (2016).

30. Han, J.-H. et al. Neuronal Competition and Selection During Memory Formation. Science 316, 457–460 (2007).

31. Josselyn, S. A. & Frankland, P. W. Memory Allocation: Mechanisms and Function. Annu. Rev. Neurosci. 41, 389–413 (2018).

32. Maristany De Las Casas, E., et al. Tuft dendrites in frontal motor cortex enable flexible learning. Preprint at 10.1126/science.adx4358 (2026).

33. Rigotti, M. et al. The importance of mixed selectivity in complex cognitive tasks. Nature 497, 585–590 (2013).

34. Crawley, D. et al. Modeling flexible behavior in childhood to adulthood shows age- dependent learning mechanisms and less optimal learning in autism in each age group. PLOS Biol. 18, e3000908 (2020).

35. Liu, Y. & Wang, X.-J. Flexible gating between subspaces in a neural network model of internally guided task switching. Nat. Commun. 15, 6497 (2024).

36. Song, H. F., Yang, G. R. & Wang, X.-J. Reward-based training of recurrent neural networks for cognitive and value-based tasks. eLife 6, e21492 (2017).

37. 37. Wierda, T., Dora, S., Pennartz, C. M. A. & Mejias, J. F. Diverse and flexible behavioral strategies arise in recurrent neural networks trained on multisensory decision making. Preprint at 10.1101/2023.10.28.564511 (2023).

38. Hattori, R. & Komiyama, T. Context-dependent persistency as a coding mechanism for robust and widely distributed value coding. Neuron 110, 502–515.e11 (2022).

39. Siniscalchi, M. J., Phoumthipphavong, V., Ali, F., Lozano, M. & Kwan, A. C. Fast and slow transitions in frontal ensemble activity during flexible sensorimotor behavior. Nat. Neurosci. 19, 1234–1242 (2016).

40. Yang, J.-H. & Kwan, A. C. Secondary motor cortex: Broadcasting and biasing animal’s decisions through long-range circuits. in International Review of Neurobiology vol. 158 443–470 (Elsevier, 2021).

41. Murakami, M., Vicente, M. I., Costa, G. M. & Mainen, Z. F. Neural antecedents of self-initiated actions in secondary motor cortex. Nat. Neurosci. 17, 1574–1582 (2014).

42. Nashaat, M. et al. The neural mechanisms of fast versus slow decision-making. Preprint at 10.1101/2024.08.22.608577 (2024).

43. Wang, T.-Y., Liu, J. & Yao, H. Control of adaptive action selection by secondary motor cortex during flexible visual categorization. eLife 9, e54474 (2020).

44. Barthas, F. & Kwan, A. C. Secondary Motor Cortex: Where ‘Sensory’ Meets ‘Motor’ in the Rodent Frontal Cortex. Trends Neurosci. 40, 181–193 (2017).

45. Hooks, B. M. et al. Organization of Cortical and Thalamic Input to Pyramidal Neurons in Mouse Motor Cortex. J. Neurosci. 33, 748–760 (2013).

46. Larkum, M. A cellular mechanism for cortical associations: an organizing principle for the cerebral cortex. Trends Neurosci. 36, 141–151 (2013).

47. Suzuki, M. & Larkum, M. E. General Anesthesia Decouples Cortical Pyramidal Neurons. Cell 180, 666–676.e13 (2020).

48. Nashaat, M. A., Oraby, H., Sachdev, R. N. S., Winter, Y. & Larkum, M. E. Air-Track: a real-world floating environment for active sensing in head-fixed mice. J. Neurophysiol. 116, 1542–1553 (2016).

49. Augusto, E. et al. Secondary motor cortex tracks decision value during the learning of a non-instructed task. Cell Rep. 44, 115152 (2025).

50. Secondary motor cortex: Broadcasting and biasing animal’s decisions through long-range circuits. in International Review of Neurobiology vol. 158 443–470 (Elsevier, 2021).

51. Feng, G. et al. Imaging Neuronal Subsets in Transgenic Mice Expressing Multiple Spectral Variants of GFP. Neuron 28, 41–51 (2000).

52. Albarran, E., Raissi, A., Jáidar, O., Shatz, C. J. & Ding, J. B. Enhancing motor learning by increasing the stability of newly formed dendritic spines in the motor cortex. Neuron 109, 3298–3311.e4 (2021).

53. Kastellakis, G., Tasciotti, S., Pandi, I. & Poirazi, P. The dendritic engram. Front. Behav. Neurosci. 17, 1212139 (2023).

54. Wang, X.-J. & Yang, G. R. A disinhibitory circuit motif and flexible information routing in the brain. Curr. Opin. Neurobiol. 49, 75–83 (2018).

55. Letzkus, J. J., Wolff, S. B. E. & Lüthi, A. Disinhibition, a Circuit Mechanism for Associative Learning and Memory. Neuron 88, 264–276 (2015).

56. Hofer, S. B., Mrsic-Flogel, T. D., Bonhoeffer, T. & Hübener, M. Experience leaves a lasting structural trace in cortical circuits. Nature 457, 313–317 (2009).

57. Hedrick, N. G., Wright, W. J. & Komiyama, T. Local and global predictors of synapse elimination during motor learning. Sci. Adv. 10, eadk0540 (2024).

58. Yasuda, M., Nagappan-Chettiar, S., Johnson-Venkatesh, E. M. & Umemori, H. An activity-dependent determinant of synapse elimination in the mammalian brain. Neuron 109, 1333–1349.e6 (2021).

59. Rangaraju, V., Lauterbach, M. & Schuman, E. M. Spatially Stable Mitochondrial Compartments Fuel Local Translation during Plasticity. Cell 176, 73–84.e15 (2019).

60. Johnson, C. M., Peckler, H., Tai, L.-H. & Wilbrecht, L. Rule learning enhances structural plasticity of long-range axons in frontal cortex. Nat. Commun. 7, 10785 (2016).

61. McCloskey, M. & Cohen, N. J. Catastrophic Interference in Connectionist Networks: The Sequential Learning Problem. in Psychology of Learning and Motivation vol. 24 109–165 (Elsevier, 1989).

62. Chavlis, S. & Poirazi, P. Dendrites endow artificial neural networks with accurate, robust and parameter-efficient learning. Nat. Commun. 16, 943 (2025).

63. Iyer, A. et al. Avoiding Catastrophe: Active Dendrites Enable Multi-Task Learning in Dynamic Environments. Front. Neurorobotics 16, 846219 (2022).

64. Holtmaat, A. et al. Long-term, high-resolution imaging in the mouse neocortex through a chronic cranial window. Nat. Protoc. 4, 1128–1144 (2009).

65. Phoumthipphavong, V., Barthas, F., Hassett, S. & Kwan, A. C. Longitudinal Effects of Ketamine on Dendritic Architecture *In Vivo* in the Mouse Medial Frontal Cortex. eneuro 3, ENEURO.0133-15.2016 (2016).

66. Pagkalos, M., Chavlis, S. & Poirazi, P. Introducing the Dendrify framework for incorporating dendrites to spiking neural networks. Nat. Commun. 14, 131 (2023).

67. Papadopoulos, P. M., Katz, M. J. & Bruno, G. NPACI Rocks: tools and techniques for easily deploying manageable Linux clusters. Concurr. Comput. Pract. Exp. 15, 707–725 (2003).

